# The two-component system CopRS maintains femtomolar levels of free copper in the periplasm of *Pseudomonas aeruginosa* using a phosphatase-based mechanism

**DOI:** 10.1101/2020.05.13.092544

**Authors:** Lorena Novoa-Aponte, Fernando C. Soncini, José M. Argüello

**Author notes:** To whom correspondence should be addressed: Dr. José M. Argüello.

## Abstract

Two component systems control periplasmic Cu^+^ homeostasis in Gram-negative bacteria. In characterized systems such as *Escherichia coli* CusRS, upon Cu^+^ binding to the periplasmic sensing domain of CusS, a cytoplasmic phosphotransfer domain phosphorylates the response regulator CusR. This drives the expression of efflux transporters, chaperones, and redox enzymes to ameliorate metal toxic effects. Here, we show that the *Pseudomonas aeruginosa* two component sensor histidine kinase CopS exhibits a Cu-dependent phosphatase activity that maintains a non-phosphorylated CopR when the periplasmic Cu levels are below its activation threshold. Upon Cu^+^ binding to the sensor, the phosphatase activity is blocked and the phosphorylated CopR activates transcription of the CopRS regulon. Supporting the model, mutagenesis experiments revealed that the Δ*copS* strain showed constitutive high expression of the CopRS regulon, lower intracellular Cu^+^ levels, and larger Cu tolerance when compared to wild type cells. The invariant phospho-acceptor residue His_235_ of CopS was not required for the phosphatase activity itself, but necessary for its Cu-dependency. To sense the metal, the periplasmic domain of CopS binds two Cu^+^ ions at its dimeric interface. Homology modeling of CopS based on CusS structure (four Ag^+^ binding sites) clearly explains the different binding stoichiometries in both systems. Interestingly, CopS binds Cu^+/2+^ with 30 × 10^−15^ M affinities, pointing to the absence of free (hydrated) Cu^+/2+^ in the periplasm.

**IMPORTANCE:** Copper is a micronutrient required as cofactor in redox enzymes. When free, copper is toxic, mismetallating proteins, and generating damaging free radicals. Consequently, copper overload is a strategy that eukaryotic cells use to combat pathogens. Bacteria have developed copper sensing transcription factors to control copper homeostasis. The cell envelope is the first compartment that has to cope with copper stress. Dedicated two component systems control the periplasmic response to metal overload. This manuscript shows that the copper sensing two component system present in Pseudomonadales exhibits a signal-dependent phosphatase activity controlling the activation of the response regulator, distinct from previously described periplasmic Cu sensors. Importantly, the data show that the sensor is activated by copper levels compatible with the absence of free copper in the cell periplasm. This emphasizes the diversity of molecular mechanisms that have evolved in various bacteria to manage the copper cellular distribution.

## INTRODUCTION

Cooper is a cellular micronutrient required for redox enzymatic functions (1, 2). However, free Cu undergoes deleterious Fenton reactions, metallates noncognate binding sites, and promotes disassemble of Fe-S centers (3, 4). Early studies in the field took advantage of Cu toxicity to identify widely distributed proteins conferring metal tolerance; namely, metal sensing transcriptional regulators and efflux transporters (1, 4–7). Recent studies have, however, started to uncover regulated distributions systems that move the metal among cellular compartments and target Cu^+^ to cognate metalloproteins while maintaining the required homeostasis (8–15). These include Cu^+^ sensing transcriptional regulators, influx and efflux transmembrane transporters, chaperones, and storage molecules. In this context, bacterial cells prevent Cu toxicity by expressing some of these molecules in response to high intracellular metal conditions. The cytoplasmic response to Cu^+^ excess has been characterized in numerous Gram-positive and Gram-negative bacteria (11, 16–19). Nevertheless, periplasmic components involved in Cu^+^ homeostasis have received much less attention. A simple consideration of the Gram-negative bacterial architecture points out that periplasmic dyshomeostasis is likely to precede the cytoplasmic response to a surge of Cu^+^ influx. Supporting this idea, mathematical simulations based on Cu^+^ uptake experiments in *Pseudomonas aeruginosa* under dyshomeostasis conditions, suggest that the periplasmic Cu^+^ overload precedes the cytoplasmic unbalance (10). Moreover, periplasmic storage molecules are likely crucial for maintaining cellular Cu^+^ allocation (10).

Cytoplasmic Cu^+^ sensing transcriptional regulators are diverse, as different bacterial species have solved Cu^+^ homeostasis using alternative strategies (1, 5, 20, 21). However, the periplasmic response appears usually regulated by similar <u>t</u>wo-<u>c</u>omponent <u>s</u>ystem (TCS) (22, 23). Although absent in *Salmonella* (6), many Enterobacteriaceae (e.g., *Escherichia coli, Klebsiella pneumoniae, etc.*) modulate periplasmic Cu^+^ stress responses via the chromosomal encoded TCS CusRS and the plasmid-borne PcoRS (24–30). Instead, CopRS monitors extra cytoplasmic Cu accumulation in *Corynebacterium glutamicum* and *Synechocystis* (31–33). CopRS is also found in Pseudomonadaceae, including *Pseudomonas syringae* (34, 35), *P. aeruginosa* (9), and *Pseudomonas fluorescens* (36, 37).

Most TCSs comprise a sensor histidine kinase (SHK) and its cognate cytoplasmic response regulator (RR). The SHK is usually a homodimeric membrane receptor with a periplasmic sensor domain flanked by two transmembrane segments (Fig. S1). The C-terminal cytoplasmic domain contains the catalytic machinery (38). SHKs are bifunctional enzymes that switch between kinase and phosphatase states in a signal-dependent manner. In the kinase mode, the SHK undergoes autophosphorylation of a conserved His residue and subsequently transfers the phosphoryl group to a conserved Asp residue of its cognate RR. In the phosphatase mode, the dephosphorylated SHK catalyzes the dephosphorylation of RR (39). Phosphorylation of the RR allosterically modifies its transcriptional activity (Fig. 1A). Ultimately, the signal-dependent balance between SHK kinase and phosphatase activities determines the RR~P levels, modulating the output response (38). In the archetypical *E. coli* CusRS TCS, Cu^+^ binding to the periplasmic loop of CusS promotes its autophosphorylation, and the subsequent phosphorylation of the transcriptional regulator CusR (Fig. 1A). Supporting this model, deletion of either the SHK CusS or the RR CusR leads to a reduced tolerance to external Cu^2+^, increased intracellular Cu^+^, and lack of transcriptional activation of regulated genes (e.g. *cusC*) (24–27).

**Fig 1.**
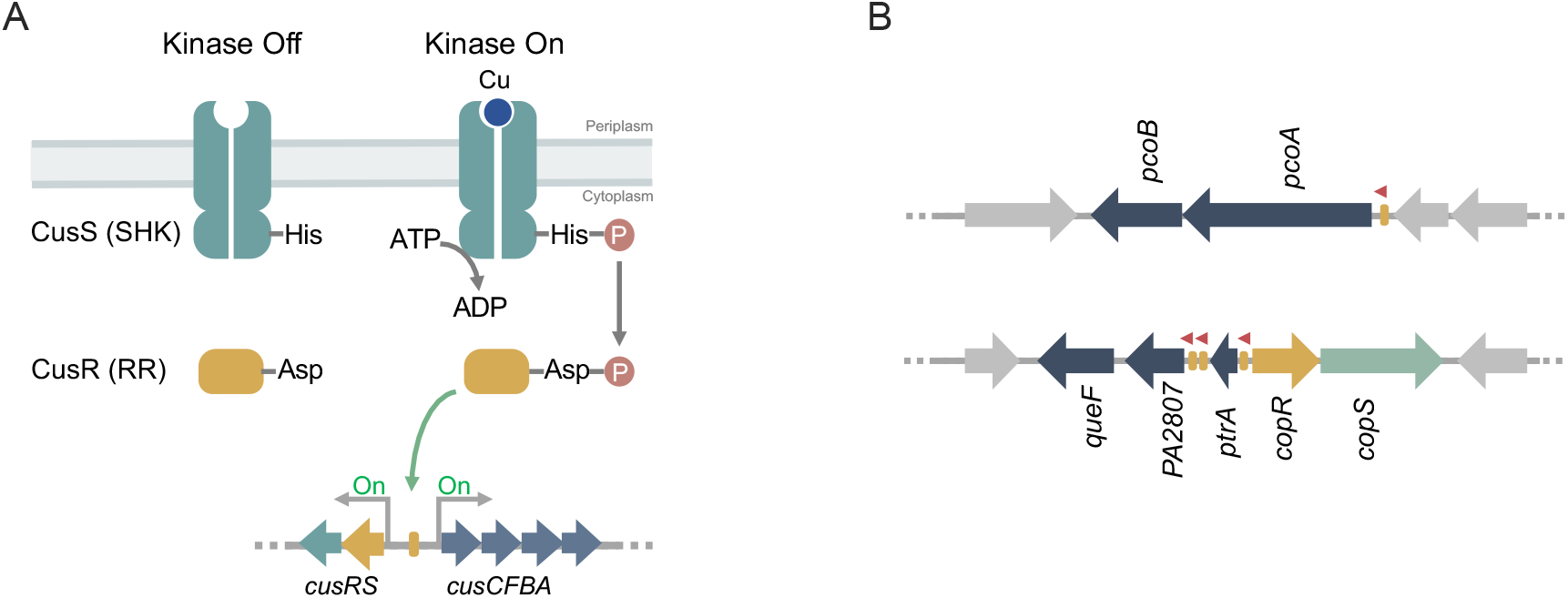
Transcriptional control mediated by TCSs. (A) Activation dynamics of canonical TCSs exemplified with the *E. coli* Cu sensing CusRS. (B) Scheme of the TCS *P. aeruginosa* CopRS regulon. Promoter regions recognized by CopR (yellow rectangles) and transcription direction (red arrows) are shown.

The regulons controlled by the canonical Cu^+^ responsive TCS are limited to genes coding for the Resistance-Nodulation-Division (RND) transporter CusCFBA (26), the PcoABCDRSE (27), and CopABCDRS systems (34, 35). However, Cu^+^ homeostatic pathways do not behave as evolutionary units. Instead, distinct species assemble different repertoires of metal handling proteins to achieve periplasmic Cu^+^ homeostasis (21). In particular, the *P. aeruginosa* CopRS regulon includes genes coding for an outer membrane transporter (PcoB), a multicopper oxidase (PcoA), and auxiliary proteins (PtrA, PA2807, QueF) whose role in periplasmic Cu^+^ distribution is still unclear (40–42) (Fig. 1B). Interesting, the *P. aeruginosa* CusCBA transporter is not part of the CopRS regulon but is rather controlled by the cytoplasmic Cu^+^ sensor CueR (9). Given the distinct architecture of the CopRS regulon, could it be expected a distinct sensing/activating mechanism for the control of periplasmic Cu^+^ homeostasis in *P. aeruginosa*?

The structure of the isolated periplasmic domain of *E. coli* CusS shows four Ag^+^ (acting as Cu^+^ analog) binding sites per dimer (43). Two sites are symmetrically located at the dimer interface, and two are situated in outer loops of separated monomers. Reported estimates of metal-sensor affinities are limited and quite different. The *E. coli* CusS interacts with Ag^+^ with an affinity in the μM range (44), while *Synechocystis* CopS binds Cu^2+^ with high sub-attomolar affinity (32). Then, significant aspects of sensor activation appear undefined. Consider that selectivity and sensitivity will determine the level of free metal in the periplasm and provide evidence for the Cu redox status. This is, can both Cu^+^/Cu^2+^ bind the sensor? What is the affinity of the sensor for these ions?

Here, we report that the transcriptional control of the CopRS regulon in *P. aeruginosa* relies on the Cu-dependent phosphatase activity of CopS, rather than on its kinase activity. The RR CopR appears constitutively phosphorylated. However, in the absence of Cu, CopS dephosphorylates CopR shutting down the transcriptional response to Cu^+^. When the periplasmic Cu^+^ level rises, the phosphatase activity of CopS is blocked, allowing the accumulation of phosphorylated CopR (CopR~P) which promotes the expression of the periplasmic Cu^+^-homeostasis network. Finally, CopS binds both Cu^+^ and Cu^2+^ with similar high affinities ensuring the absence of free Cu in the periplasm.

## RESULTS

CopRS controls *P. aeruginosa* periplasmic Cu^+^ homeostasis (9). Notably, there are significant differences between the CopRS regulon and those of other characterized Cu^+^ sensing TCSs, e.g. *E. coli* CusRS. The likely presence of additional mechanistic and molecular differences warranted a closer examination of CopRS function.

### Mutation of CopS leads to Cu tolerance

We initiated our studies by looking at the growth rate of Δ*copS* and Δ*copR* mutant strains in the presence of external metal. Based on the mechanism of described TCS (Fig. 1A), it was expected that the lack of either CopS or CopR would lower the cellular tolerance to external Cu^2+^. As anticipated, the Δ*copR* strain was more susceptible to Cu^2+^ than the WT strain (Fig. 2). In contrast, two independent *copS* transposon mutants, PW5705 and PW5706 (Fig. S1), were surprisingly much more tolerant to external Cu^2+^ than the WT strain. As these phenotypes were reversed by complementation with the corresponding genes all subsequent experiments were performed with the Δ*copS* PW5706 strain. For comparison, in addition to the WT strain, the well characterized Cu^+^ sensitive Δ*copA1* mutant strain was also included as control in this initial phenotypical characterization (8).

**Fig 2.**
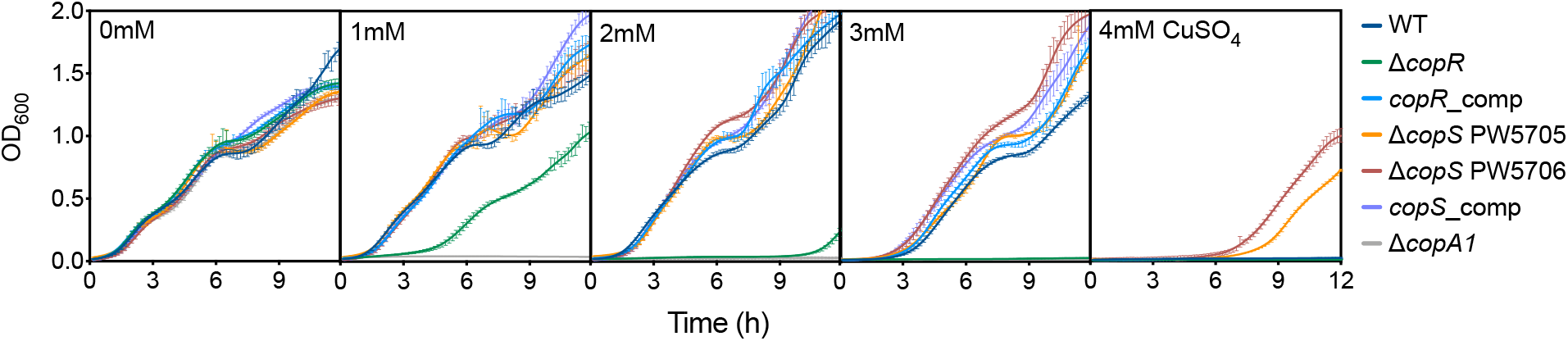
Cu tolerance of Δ*copR* and Δ*copS* mutant strains. Growth rate of WT, Δ*copR*, Δ*copS* (PW5705 and PW5706), Δ*copA1*, and the CopR and CopS complemented strains in the presence of 0 - 4 mM CuSO_4_. Data are the mean ± SEM of at least three independent experiments.

Importantly, these growth phenotypes were the consequence of significantly different levels of intracellular Cu^+^ upon exposure to CuSO_4_ (Fig. 3). Thus, the Δ*copR* mutant strain accumulated more Cu^+^, while the Δ*copS* cells stored less metal than the WT strain. Again, alterations in Cu^+^ levels were reversed by gene complementation of the mutant strains. These differences in Cu tolerance and cellular metal levels observed for the Δ*copR* and Δ*copS* mutant strains cannot be explained by the currently accepted model for the *E. coli* TCS CusRS (Fig. 1A) and suggest an alternative mechanism for coupling periplasmic Cu^+^ sensing and gene expression in *P. aeruginosa*.

**Fig 3.**
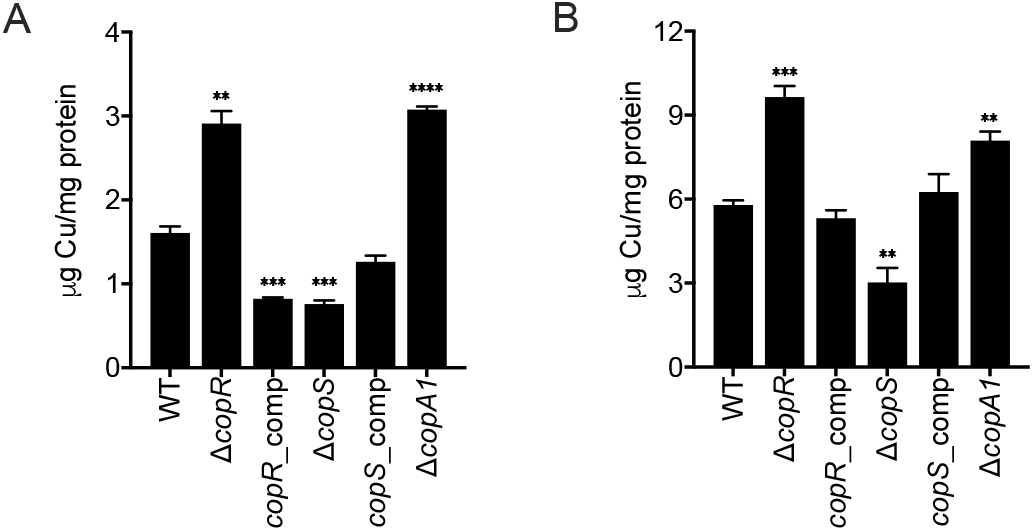
Whole cell Cu levels in WT, Δ*copR*, Δ*copS*, Δ*copA1*, and CopR and CopS complemented strains after 10 min exposure to (A) 2 mM CuSO_4_ and (B) 4 mM CuSO_4_. Data are the mean ± SEM of three independent experiments. Significant differences from values with the WT strain as determined by unpaired two-tailed Student’s t-test are \*\**P* < 0.01, \*\*\**P* < 0.001, \*\*\*\**P* < 0.0001.

### The CopRS regulon is constitutively expressed in the ΔcopS mutant strain independently of Cu^+^ levels

Toward understanding the increased Cu tolerance and intracellular levels in the Δ*copS* strain, we investigated the transcriptional response to Cu^2+^ exposure of the CopRS regulon in the Δ*copR* and Δ*copS* mutant strains. We have described that CopRS controls the expression of *pcoA*, *pcoB*, *ptrA*, *queF,* and *PA2807* coding for periplasmic and outer membrane proteins (Fig. 1B) (9). As previously observed in the WT strain, genes of the CopRS regulon are induced in response to external Cu^2+^ exposure (Fig. 4). As expected, their Cu-induced expression was abolished in the Δ*copR* mutant. Quite the opposite, the Δ*copS* mutant strain showed a constitutive activation of all the genes of the CopRS regulon, even in the absence of the Cu^2+^ stimulus. In the Δ*copS* background, high expression levels of these genes were similar either under the absence of added Cu^2+^, low, non-deleterious 0.5 mM Cu^2+^ levels, intermedium toxic 2 mM Cu^2+^, and high lethal 4 mM Cu^2+^ (Fig. S2). This suggests that CopS is not necessary to activate, i.e. phosphorylate, CopR. Furthermore, it implies that CopR is phosphorylated independently of CopS. The activation of CopR in the Δ*copS* mutant in the absence of supplemented Cu^2+^ points to a mechanism where the phosphatase activity of CopS maintains low levels of CopR~P under noninducing conditions. This defect in the Δ*copS* strain to maintain the system *off* in absence of Cu^+^ was reversed in the complemented strain (Fig. 4). The transcriptional analysis also showed that the expression of the *copRS* operon is not autoregulated by CopRS (Fig. S3). This is, even though *copRS* expression is induced in response to Cu^+^, no defects on its expression were observed in the Δ*copR* or the Δ*copS* mutant strains. Noticeably, the repressed transcription of *oprC*, codding for the outer membrane Cu importer (9, 45), was further repressed on the Δ*copS* mutant strain, consistent with the Cu^+^ tolerant phenotype exhibited by this strain (Fig. S4A). Conversely, the increased transcription of genes in the CueR regulon (*copA1* and *cusA*) in response to Cu^+^, was not altered neither in Δ*copR* nor Δ*copS* mutant strains (Fig. S4B). This confirms that the lack of transcriptional control observed in the Δ*copR* and Δ*copS* mutant strains is limited to genes of the CopRS regulon.

**Fig 4.**
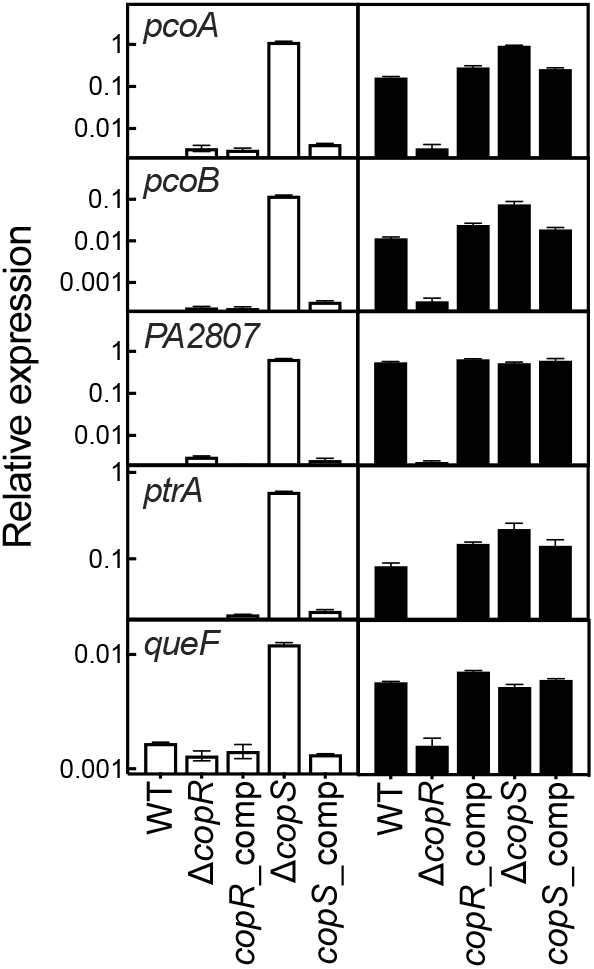
Expression of genes in the CopRS regulon in WT, Δ*copR,* Δ*copS* and the corresponding complemented strains in the absence (white) and the presence (black) of 0.5 mM CuSO_4_ (5 min treatment). Transcript levels of *pcoA*, *pcoB*, *PA2807*, *ptrA,* and *queF* genes are plotted relative to that of the housekeeping gene *PA4268*. Data are the mean ± SEM of three independent experiments.

### His_235_ acts as a switch to turn on/off the CopS signaling pathway

The cytoplasmic region of the SHK sensory proteins contains the catalytic phosphotransfer domain able to switch between kinase and phosphatase activities in a signal-dependent manner (39, 46). In the canonical SHKs, this phosphotransfer domain contains an invariant His that autophosphorylates in the first step of the signaling cascade, activating the kinase-state of the SHK. Subsequently, the RR protein is phosphorylated in a highly conserved phospho-acceptor Asp leading to transcription activation (47) (Fig. 1A). In contrast to the kinase state, in the phosphatase state a dephosphorylated SHK removes the phosphate group from the RR~P (39). The kinase and phosphatase states are mutually exclusive. In some cases, the activation of the kinase state is associated with phosphatase deactivation with the consequent accumulation of RR~P. The observed phenotypes in Δ*copS* and Δ*copR* strains suggest that in the absence of Cu^+^, CopS acts as a phosphatase dephosphorylating CopR. Then, when CopS senses Cu^+^, its phosphatase would be inactivated leading to a rise of CopR~P, triggering the expression of the CopRS regulon. Testing these ideas, the phosphorylatable residues, His_235_ in CopS and Asp_51_ in CopR, were identified by sequence alignment with characterized TCS (Fig. S5). Site-directed mutagenesis was performed to generate Asp_51_Ala and Asp_51_Glu replacements in CopR and His_235_Ala in CopS coding sequences. The resulting constructs were employed to complement the corresponding Δ*copR* and Δ*copS* mutant strains.

Fig. 5A shows that, as expected, replacements Asp_51_Ala and Asp_51_Glu in CopR lead to growth phenotypes comparable to that of Δ*copR* strain. Conversely, the His_235_Ala CopS mutant was able to revert the Δ*copS* Cu tolerance phenotype (Fig. 5B), suggesting that the phosphatase remains active and, as the His was replaced by Ala, His_235_ would not be required for this activity. Analysis of the transcriptional activation of genes in the CopRS regulon further supports this idea. Fig. 6 shows the regulation of *pcoB* in the relevant mutant strains. Again, the mutant CopR proteins were unable to activate *pcoB* expression in the presence of external Cu^2+^, a lack of function associated with the absence of phosphorylation of Asp_51_. In absence of supplemented Cu^2+^, *pcoB* transcription remained low in His_235_Ala CopS mutant, similar to the level observed in WT strain and contrary to the increased expression in the Δ*copS* mutant strain. Conversely, in the presence of Cu^2+^, the His_235_Ala mutant did not behave as the WT nor as the Δ*copS* mutant, but as the Δ*copR* mutant. The lack of transcriptional activation suggests that the His_235_Ala mutant was insensitive to Cu^+^, in other words this mutation locked CopS in a phosphatase-ON state regardless of the presence of metal.

**Fig 5.**
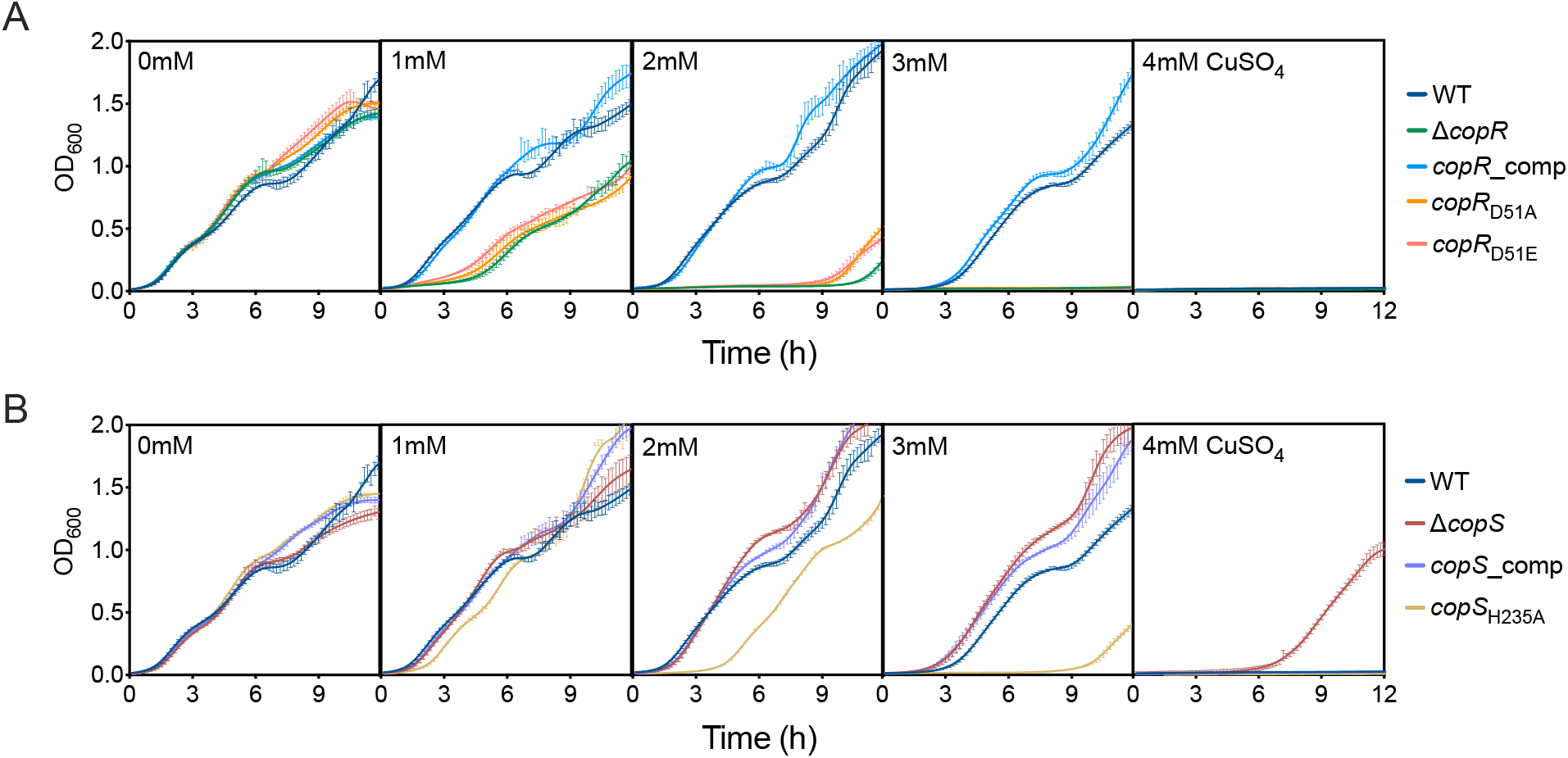
Cu tolerance of Δ*copR* and Δ*copS* mutant strains complemented with CopR and CopS lacking the phosphorylatable residues. (A) Growth rate of the Δ*copR* mutant complemented with *copR* coding for substitutions Asp_51_Ala and Asp_51_Glu in the presence of 0 - 4 mM CuSO_4_. (B) Growth rate of the Δ*copS* mutant complemented with the *copS* gene coding for substitution His_235_Ala in the presence of 0 - 4 mM CuSO_4_. Data are the mean ± SEM of three independent experiments.

**Fig 6.**
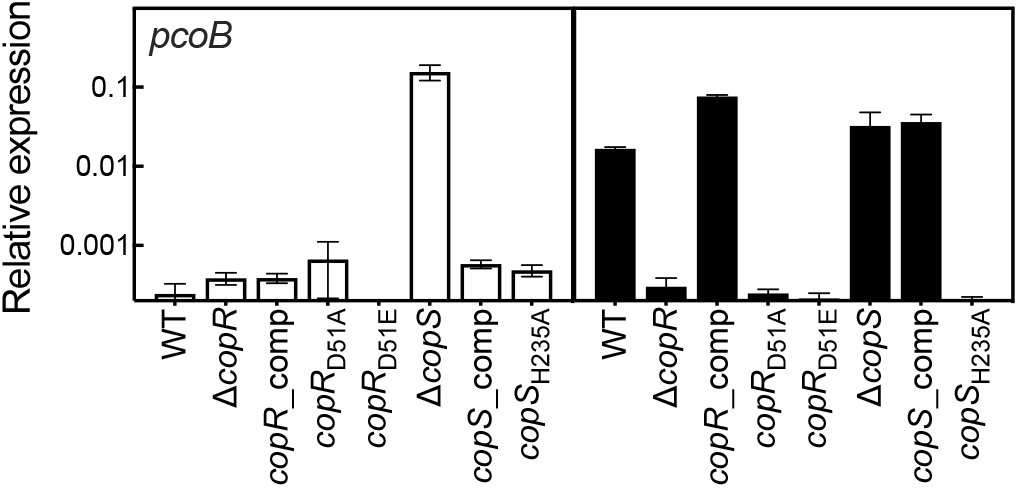
Expression of *pcoB* in Δ*copR* and Δ*copS* mutant strains complemented with CopR and CopS lacking the phosphorylatable residues in the absence (white) and the presence (black) of 2 mM CuSO_4_ (5 min treatment). Δ*copR* mutant was complemented with *copR* coding for substitutions Asp_51_Ala and Asp_51_Glu. Δ*copS* mutant was complemented with the *copS* gene coding for substitution His_235_Ala. Transcript levels of *pcoB* are plotted relative to the housekeeping gene *PA4268*. Data are the mean ± SEM of three independent experiments.

### CopS periplasmic sensor domain binds two Cu^+^ ions per functional unit

Most TCS sensors are homodimeric membrane receptors. The periplasmic sensor domain of CopS, flanked by two transmembrane segments (Fig. S1), extends between residues 34 and 151 (CopS_(34-151)_). The function of the system relies on its ability to bind cognate metal ions. To explore CopS metal binding properties, the *P. aeruginosa* CopS_(34-151)_ sensor domain was heterologously expressed and purified to homogeneity (Fig. S6). The isolated domain bound 2.3 ± 0.5 Cu^+^ ions per CopS_(34-151)_ dimer. This differs from the four Ag^+^ (used as Cu^+^ analog) per dimer stoichiometry observed in *E. coli* CusS (43). However, the periplasmic sensor domain of CopS homolog proteins is considerably shorter than the CusS domain, lacking a loop containing residues (Ser_84_, Met_133_, Met_135_ and His_145_) involved in metal biding in CusS (Fig. S5B). In effect, a phylogenetic tree built with sequences homologues to CopS and CusS (>45% identity) shows a clear evolution of two distinct subgroups of CusS homologues in Enterobacterales and in Burkholderiales, and a separate group of CopS homologues in Pseudomonadales (Fig. S7). This structural differences leading to the alternative stoichiometry can be more easily observed when the homology modeling of *P. aeruginosa* CopS is overlapped with the crystal structure of the Ag^+^-bound periplasmic sensor domain of *E. coli* CusS (43) (Fig. 7). The two symmetric metal binding sites fully conserved in both, CopS and CusS, are located at the dimeric interface. Each site is formed by two invariant His residues (His_41_ and His_140_ in CopS), one from each dimer subunit. A Phe likely interacting with the metal in CusS, is also conserved in CopS (Phe_42_). These are probably the Cu^+^ sensing sites involved in signal transduction. On the other hand, the structural comparison clearly shows that the loop containing the additional metal binding sites of CusS is missing in CopS (orange loops, Fig. 7).

**Fig 7.**
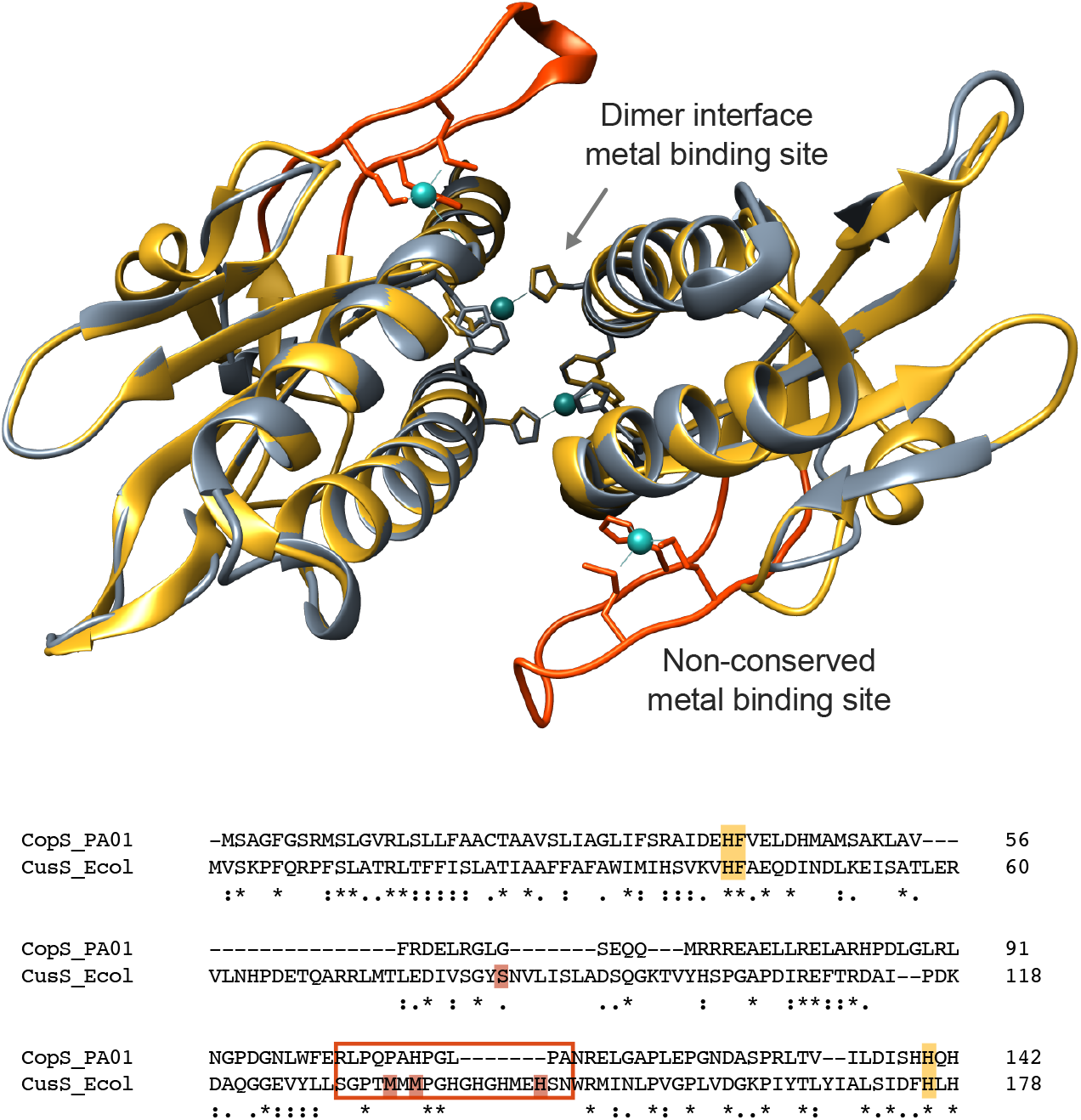
Structural superposition of the periplasmic Cu^+^ binding loop of *P. aeruginosa* CopS (gray) and *E. coli* CusS (yellow). The structure of CopS was modelled using the CusS structure as template (PDB ID: 5KU5 (43)). Conserved Cu binding sites at the dimeric interface (His_41_, Phe_42_, and His_140_) are shown as sticks in the structural model and highlighted in yellow in the sequence alignment. The Cu^+^ binding sites within the CusS orange loops (framed in rectangle in the alignment), are not conserved in CopS.

### The CopS periplasmic sensor binds Cu^+/2+^ with femtomolar affinities

By analogy on how cytoplasmic sensors metal affinities are tuned to maintain free metal levels (48, 49), the affinity of CopS for Cu^+^ ions will certainly have determinant effects on free (hydrated) Cu^+^ ions levels in the periplasm. Exploring the binding of Cu^+^ to CopS, we measured the sensor metal binding affinity using competing ligands. The ligands were present in excess to ensure effective competition. In all cases, the determinations were performed assuming that both Cu sites at the CopS dimer interface were functionally independent and thermodynamically indistinguishable. Initial determinations of CopS_(34-151)_ affinity for Cu^+^ using the bathocuproine disulfonate (BCS) as competitor, showed a limited but measurable decrease in the absorbance of the [Cu^I^(BCS)_2_^3−^] complex, corresponding to a *K_D_* value of CopS_(34-151)_ for Cu^+^ of 22 × 10^−15^ M. However, it was apparent that CopS was not an effective competitor with BCS for the metal. Instead 2,2′-bicinchoninic acid (BCA), with a lower affinity for copper compared to BCS, appeared more appropriate to measure affinities in the femtomolar range (50). Using BCA as the competing ligand and fitting titration curves to Eq. 2, a CopS_(34-151)_-Cu^+^ *K_D_* = 27.7 ± 0.7 × 10^−15^ M was obtained (Fig. 8A). This appears within the range of affinities observed for many other Cu^+^ binding molecules (11, 50, 51).

**Fig 8.**
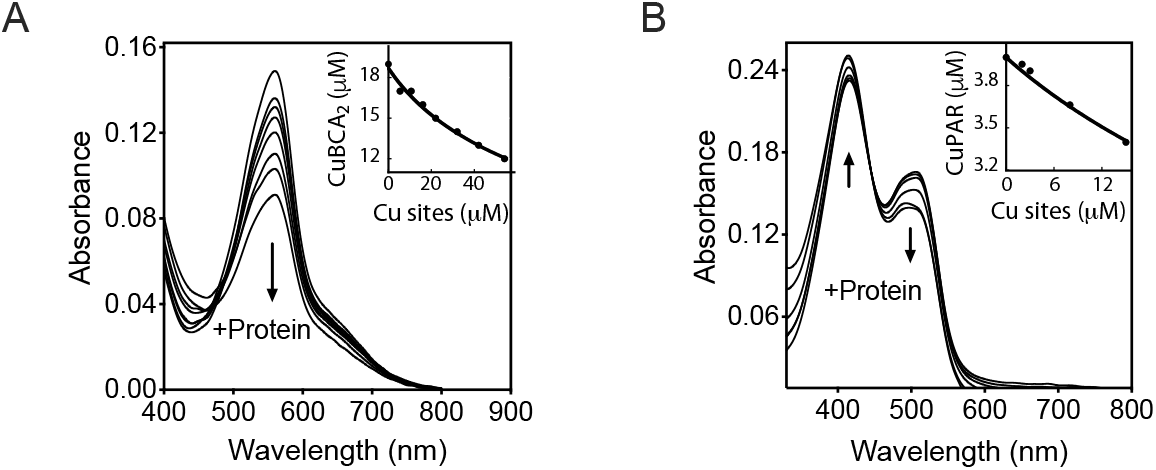
Determination of the dissociation constants *K_D_* of the periplasmic Cu binding loop of CopS_(34-151)_. (A) Spectrophotometric titration of 100 μM BCA and 18.7 μM Cu^+^ with increasing concentrations of CopS_(34-151)_. The arrow indicates the decrease in absorbance at 562 nm upon protein addition. The inset shows the fitting of the data set to equation (2) with a *K_D_* = 27.7 ± 0.7 × 10^−15^ M (R^2^ 0.992). Two Cu sites per CopS dimer are assumed. (B) Spectrophotometric titration of 10 μM PAR and 4 μM Cu^2+^ with increasing concentrations of CopS_(34-151)_. The arrows indicate the increase in absorbance at 415 nm and the decrease at 562 nm upon protein addition. The inset shows the fitting of the data set to equation (4) with a *K_D_* = 33.2 ± 1.5 × 10^−15^ M (R^2^ 0.984).

On the other hand, *Synechocystis* CopS binds Cu^2+^ with high sub-attomolar affinity (Cu^+^ binding stoichiometry was not reported) (32). Exploring the possibility of high affinity Cu^2+^ binding to *P. aeruginosa* CopS, the chromogenic ligand 4-(2-pyridylazo)resorcinol (PAR) was used as competitive ligand with a purified Step-tagged CopS_(34-151_ (Fig. S6). A CopS_(34-151)_-Cu^2+^ *K_D_* = 33.2 ± 1.5 × 10^−15^ M was observed (Fig.8B). Consequently, it is apparent that CopS_(34-151)_ binds both, Cu^+^ and Cu^2+^, with quite similar affinities in the femtomolar range. These high affinities provide insights into *in vivo* steady state and virtual absence of free Cu ions in the bacterial periplasm.

## DISCUSSION

The relevance of the periplasmic Cu pool in the *P. aeruginosa* response to Cu stress is well established (10, 52). Results presented here show novel important characteristic of the *P. aeruginosa* TCS CopRS. The sensor has a negative control mechanism based on its phosphatase rather than kinase activity. At the dimer interface, it binds two Cu^+/2+^ ions with femtomolar affinities, likely resulting in the absence of periplasmic free Cu. CopRS unique mechanism is in line with the other distinct features of *P. aeruginosa* Cu homeostasis, namely cytoplasmic and periplasmic sensors with singular regulons, a RND-transporter regulated by the cytoplasmic sensor, and multiple cytoplasmic Cu^+^ chaperones and efflux P_1B_-ATPases (8–11, 53). The emerging model challenges a number of ideas associated with the early study of *E. coli* CusRS TCS. Along with *Salmonella,* that has a distinct Cu^+^ balance mechanisms (6), *P. aeruginosa* provides a clear example of alternative approaches to achieve Cu homeostasis.

### CopS Cu-controlled phosphatase activity mediates signal transduction

Characterization of CopRS was initiated by analyzing the tolerance of Δ*copS* and Δ*copR* strains to external Cu^2+^. While an increased sensitivity was expected based on the reported phenotypes of *E. coli* Δ*cusS* and Δc*usR* strains, the Δ*copS* strain showed higher tolerance to the external metal. Although unexpected, this phenomenon has been previously observed albeit unnoticed. It was reported that deletion of *P. aeruginosa* CopS did not compromise the ability of the bacteria to grow in the presence of Cu^2+^ (54). Furthermore, while there was no evident Cu-induced expression of a *lacZ* transcriptional fusion to a *Pseudomonas putida* CinRS (a CopRS ortholog) dependent promoter in a *P. aeruginosa* Δ*copR* background, constitutive Cu-independent expression of the same reporter was attained in the *P. aeruginosa* Δ*copS* background (55). Also similar to *P. aeruginosa* Δ*copS* strain, a *P. fluorescens* Δ*copS* strain was more tolerant to external Cu^2+^ (36).

The Cu resistance phenotype of the *P. aeruginosa* Δ*copS* strain, is supported by the constitutive expression of the CopRS regulon and the consequent reduced whole-cell Cu^+^ content. The simplest explanation for these observations is a mechanism where in absence of Cu^+^, the CopS phosphatase activity abrogates the induction of the CopRS regulon by maintaining low levels of CopR~P (Fig. 9). When CopS detects periplasmic Cu overload, its phosphatase activity is blocked allowing the accumulation of CopR~P, which promotes the expression of the periplasmic Cu-homeostasis network.

**Fig 9.**
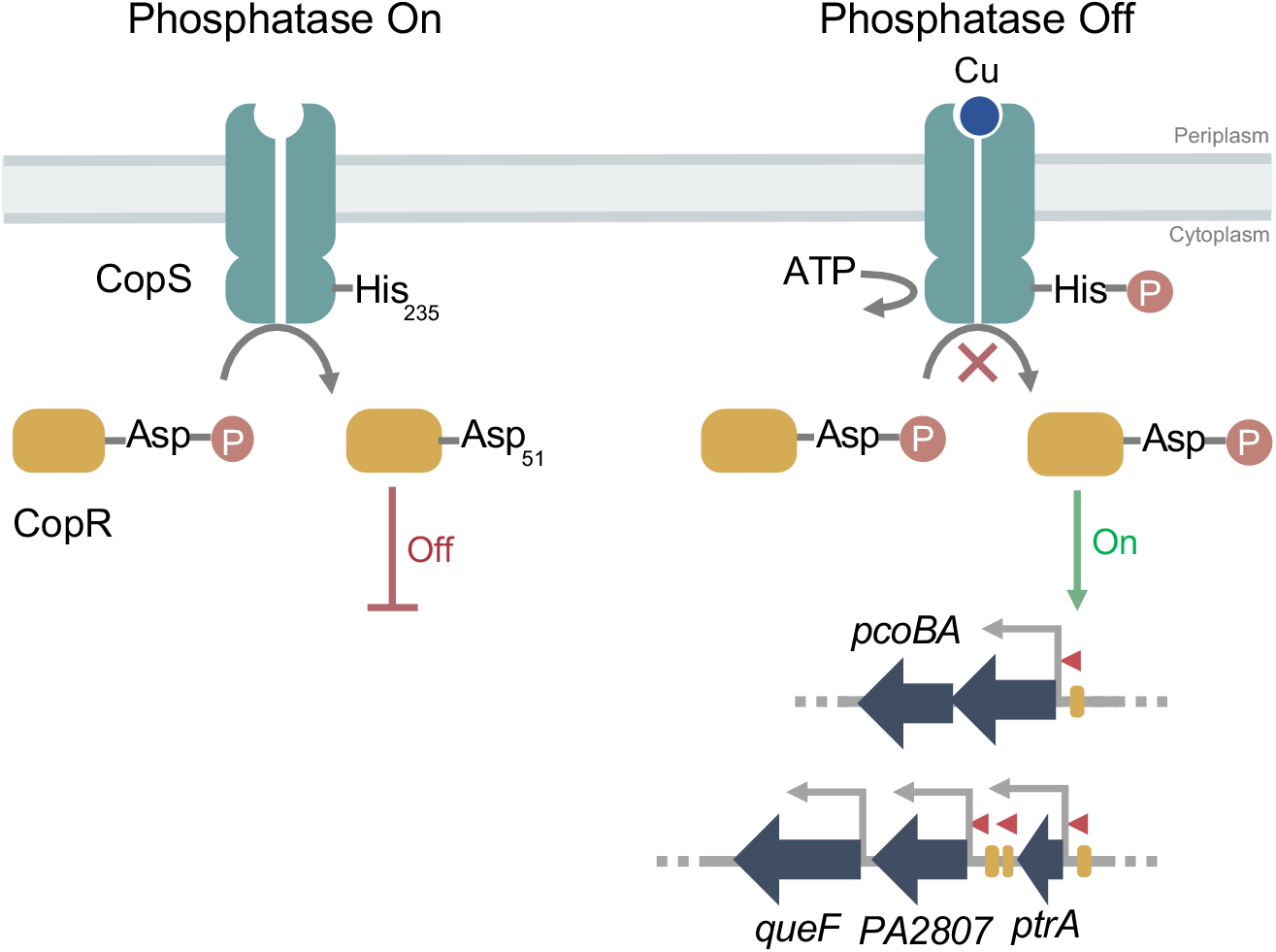
Model of the phosphatase-based mechanism of the *P. aeruginosa* CopRS. Phosphatase On: When periplasmic free Cu remains under sub-femtomolar level, the CopS phosphatase activity maintains low levels of constitutively phosphorylated CopR, shutting *off* the transcriptional response to high periplasmic Cu. Phosphatase Off: Upon Cu binding CopS autophosphorylates at His_235_. This turns *off* the CopS phosphatase activity, allowing the accumulation of phosphorylated CopR, and triggering the expression of the CopRS regulon (i.e. *pcoA*, *pcoB*, *queF*, *PA2807*, and *ptrA*).

Signal transduction by archetypical TCSs relies on bifunctional kinase/phosphatase SHKs (56). A positive action results from sensor autokinase activity and phosphotransfer to the RR; while, negative regulation involves the sensor phosphatase activity (46). The ultimate determining factor of the cascade activation is the phosphorylation status of the RR. Accumulation of a RR~P is the consequence of a signal-dependent stimulation of the sensor-kinase activity or a signal-dependent blockage of the sensor-phosphatase activity. Our data suggests that CopS harbors autokinase and phosphatase activities. The signal-independent activation of the CopRS regulon in the Δ*copS* background evidences the requirement of the CopS phosphatase activity to maintain low levels of CopR~P in the absence of Cu. It is also apparent that CopS is not required for the phosphorylation of CopR, implying that an alternative mechanism for the phosphorylation of CopR should exist. There is extensive evidence that RRs can be phosphorylated by endogenous phosphor-donors like acetyl phosphate (57). Alternative mechanisms for RR phosphorylation (cross-phosphorylation) known as many-to-one or one-to-many, where many SHKs phosphorylate a given RR or a single SHK phosphorylates multiple RRs have been proposed (38, 56). It could then be argued that CopR phosphorylation might be the consequence of an unspecific crosstalk with a non-cognate SHK that occurs only in the absence of CopS. However, such crosstalk has been only observed when both, the reciprocal RR and the cognate SHK, were absent (58). These conditions are distinct from those in our experiments.

The evidence indicates that Cu-dependent CopS autokinase activity is required for the inhibition of the CopS-phosphatase activity. Replacement His_235_Ala, leads to constitutive phosphatase activity regardless the periplasmic Cu^+^ levels. While this points out that His_235_ is not required for the CopS phosphatase activity, it implies that Cu-dependent CopS autophosphorylation turns *off* the CopS phosphatase activity, leading to accumulation of CopR~P. This is, as described, the dephosphorylated SHKs has phosphatase activity (39, 46).

### CopS binds two Cu^+/2+^ with femtomolar affinities

Its Cu binding characteristics is what defines the function of CopS. We determined that *P. aeruginosa* CopS binds two metal ions with a high affinity in the 30 × 10^−15^ M range. Little information is available regarding the binding stoichiometry and affinities of Cu sensing TCS. The structure of *E. coli* CusS clearly supports a stoichiometry of four metal atoms per CusS sensing dimer (43). Two of these ions bind at the dimer interface, while the other two attach to external loops, one in each subunit. Structural comparison of *P. aeruginosa* CopS and *E. coli* CusS shows that both types of sensors would bind and sense the metal with conserved His residues at the dimer interface. However, the CusS extra sites are not conserved in CopS nor in its homologs. Regarding binding affinities, *E. coli* CusS binds Ag^+^ with a reported 8 μM affinity, measured in equilibrium dialysis experiments (44); in contrast, *Synechocystis* CopS binds Cu^2+^ with sub-attomolar affinity (32). It would be quite speculative to compare such dissimilar determinations. However, it might be instructive to consider the observed 10^−19^–10^−21^ M affinities of cytoplasmic copper sensors in general (51, 59); and those determined for the cytoplasmic triad CopZ2/CueR/CopZ1 of *P. aeruginosa*, with relative affinities for Cu^+^ ranging between 10^−15^ – 10^−17^ M (9, 11). The weaker affinity of CopS compared to the cytoplasmic regulators and chaperones, is likely the consequence of a metal binding site formed by His rather than Cys residues. This is a logic arrangement, given the possible oxidation of proximal Cys under periplasmic redox stress. Importantly, a femtomolar affinity still supports the idea that there would not be free Cu^+/2+^ in the cell periplasm, as shown for the cytoplasm (51, 60). However, the relative binding strength of CopS is likely to be linked to those of periplasmic Cu^+^ chaperones that exchange metal with the sensor. This is, the proteins should be able to exchange the metal. Although, as shown with cytoplasmic chaperone/sensor partners, the protein-protein binding affinity will have a significant effect in the final exchange constant (11).

CopS binds both, Cu^+^ and Cu^2+^, with similar high affinity. It is accepted that cytoplasmic transporters and chaperones, bind and distribute Cu^+^. However, the periplasm is a more oxidizing compartment (61, 62). PcoA is a multicopper oxidase present in the *P. aeruginosa* periplasm (63). It has been proposed that periplasmic enzymes might catalyze Cu^+^ oxidation to the assumed less toxic Cu^2+^ (64). However, free (hydrated) Cu^+^ would be spontaneously oxidized by O_2_ in an aerobic environment. Then, the redox status of periplasmic Cu is unclear. Beyond the goals of this report, we presume that Cu oxidation state will depend on the molecule interacting and delivering Cu to CopS. In any case, the capability to bind Cu^+/2+^ might serve CopS to sense the metal under redox stress.

### The distinct CopRS mechanism is in line with the singular architecture of *P. aeruginosa* Cu homeostasis system

*E. coli* and *Salmonella* are the frequent models to explore transition metal homeostasis in Gram-negative bacteria. However, recent studies of *P. aeruginosa* have begun to show different novel molecular strategies to sense, buffer and distribute Cu^+^ (8–10, 65). For instance, consider how the regulons of both compartmental sensors, CopRS and CueR, differ among these three organisms (6, 9, 24, 66, 67). Also, analyze the multiple functionally distinct homologous Cu^+^ ATPases present in *Salmonella* and *Pseudomonas* and how these three Gram-negative bacteria have solved cytoplasmic Cu^+^-chaperoning using alternative strategies (6, 11, 68). Along these observations, the relevance of periplasmic Cu^+^ sensing, storage and transport has become more apparent. Then, it is not surprising that these model systems solve periplasmic Cu^+^ sensing either via a kinase sensor (CusRS, *E. coli*), an integration of a cytoplasmic Cu sensor with a general envelope stress response TCS (CueR-CpxRS, *Salmonella* (69)) or a phosphatase sensor (CopRS, *P. aeruginosa*). The evolutive and ecological advantages of these systems are still to be discovered and will be the subject of future enquires in the field.

## MATERIALS AND METHODS

### Bacterial strains, plasmids and growth conditions

Bacterial strains, plasmids, and primers used in this study are listed in Table S1. *P. aeruginosa* PA01 served as WT strain. Mutant strains PW5704 (Δ*copR*), PW5705 (Δ*copS*), and PW5706 (Δ*copS*) were obtained from the *P. aeruginosa* PAO1 transposon mutant library (University of Washington, Seattle, WA) (70, 71). *P. aeruginosa* strains were grown at 37°C in Luria-Bertani (LB) medium supplemented 25 μg/mL irgasan, 30 μg/mL tetracycline (mutant strains) or 30 μg/mL gentamicin (complemented strains). *E. coli* strains were grown at 37°C in LB medium supplemented with 100 μg/mL ampicillin, 30 μg/mL kanamycin, or 10 μg/mL gentamicin, depending on the plasmid selection.

### Construction of *P. aeruginosa* complemented strains

Mutant strains were complemented with the corresponding gene under control of the native promoter using the mini-Tn7 insertion system (72). Briefly, the genes and their 500 bp upstream promoter regions were amplified by PCR. The 3’ primer included a His_6_-tag coding sequence. Amplicons were cloned into the pUC18-mini-Tn7-Gm suicide delivery vector. These plasmids were used as template to introduce mutations coding for single substitutions *copR*_D51A_, *copR*_D51E_ and *copS*_H235A_ using Gibson assembly (73). The resulting plasmids were then introduced into recipient strains by conjugation, using the helper strains SM10(λpir)/pTNS2 and HB101/pRK2013. Conjugants were selected on 30 μg/mL gentamicin, 25 μg/mL irgasan, LB plates. Complemented strains were verified by PCR.

### Cu^2+^ sensitivity assay

Overnight cultures were diluted in 25 μg/mL irgasan, LB medium, adjusted to 0.05 OD_600_, and supplemented with the indicated CuSO_4_ concentration. Cell growth in liquid media was monitored for 24 h (OD_600_) at 37°C with continuous shaking using an Epoch 2 Microplate Spectrophotometer (BioTek).

### Whole cell Cu content

Cells (mid-log phase) were incubated in LB media supplemented with 0.5, 2, or 4 mM CuSO_4_. Aliquots were taken after 10 min, treated with two times molar excesses of DTT and BCS, and harvested by centrifugation at 17,000 x g, 1 min. Pellets were washed twice with 150 mM NaCl and mineralized with fuming HNO_3_ (trace metal grade) for 60 min at 80°C, and 2 M H_2_O_2_ for 60 min at room temperature. Cu levels were measured using atomic absorption spectroscopy (AAS) as described (9).

### Gene expression analysis

Cells (mid-log phase) were incubated in antibiotic-free LB media supplemented with 0.5, 2, or 4 mM CuSO_4_. 0.5 mL aliquots were taken at various times, stabilized with RNA protect Bacteria Reagent (Qiagen), and RNA isolated with RNeasy Mini Kit (Qiagen). RNA was treated with DNase I, purified by phenol/chloroform extraction and ethanol precipitated. 1 μg of RNA was used for cDNA synthesis using the ProtoScript® II kit (New England BioLabs). qPCR reactions were carried out with FastStart Essential DNA Green Master (Roche) in 10 μL final volume, using 0.25 μM of each primer (Table S1). The efficiency of primer sets was evaluated by qPCR in serial dilutions of WT cDNA. Results were normalized to 30S ribosomal protein S12 (*PA4268*) (8).

### Protein expression and purification

The *DNA fragment encoding* for the periplasmic copper binding loop of CopS_(34-151)_ was amplified from genomic DNA using 3’-end primers that introduced sequences coding either a Strep-tag or a His_6_-tag joined by a TEV cleavage site (Table S1). Resulting amplicons were cloned in the pBAD-topo vector (Invitrogen) and expressed in *E. coli* LMG194 cells. *His-tagged* CopS_(34-151)_ *was purified using Ni-NTA columns (Roche)(11). Strep-tagged* CopS_(34-151)_ *was affinity purified using Strep-Tactin^®^XT Superflow^®^ columns (IBA) (11).* Purified proteins were stored in 20% glycerol, 25 mM Tris (pH 8), 100 mM sucrose, 150 mM NaCl at −80°C. Protein concentrations were determined in accordance to Bradford (74) and purity was estimated by SDS-PAGE followed by Coomassie brilliant blue staining (Fig. S6). Proteins were purified as ≥ 90% *apo* forms as confirmed by AAS.

### Copper Binding determinations

CopS_(34-151)_-Cu^+^ binding stoichiometry was determined by incubating CopS_(34-151)_ His-tagged protein with five times molar excess of CuSO_4_ in 25 mM Hepes pH 8, 150 mM NaCl, 0.5 mM DTT for 10 min at room temperature with gentle agitation. DTT was included to reduce Cu^2+^ to Cu^+^ and prevent protein precipitation that occurs upon addition of excess Cu^+^. Unbound Cu^+^ was removed by passage through a Sephadex G-10 column (GE Healthcare) followed by two washing steps using a 3 kDa Centricon. The amount of Cu^+^ bound to protein was determined by AAS.

CopS_(34-151)_ - Cu^+^ dissociation constants (*K_D_*) were determined by competition assays with the chromogenic ligands BCS ([Cu^I^(BCS)_2_]^3−^ *β_2_’* formation constant 10^20.8^ M^−2^, *ε*_483 nm_ 13000 M^−1^ cm^−1^) and BCA ([Cu^I^(BCA)_2_]^3−^ *β_2_’* formation constant 10^17.7^ M^−2^, *ε*_562 nm_ 7900 M^−^ 1cm^−1^ (75)). Cu^+^ solutions were generated from CuSO_4_ in the presence of large excess ascorbate and NaCl which stabilizes Cu^+^ as [Cu^I^Cl_n_]^(1−n)−^ (76). Briefly, for BCS competitions, 10 μM Cu^+^, 25 μM BCS in buffer 25 mM Hepes pH 8, 150 mM NaCl, 10 mM ascorbic acid were titrated with 10 - 50 μM His-tagged CopS_(34-151)_, incubated 5 min at room temperature, and the 300-800 nm absorption spectra recorded. The same protocol was used for BCA competitions using 18.7 μM Cu^+^, 100 μM BCA and 5 - 50 μM protein instead. CopS_(34-151)_ - Cu^+^ *K_D_* were calculated by curve-fitting of the experimental data to the equilibrium equations (1) and (2) (50).

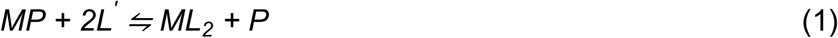

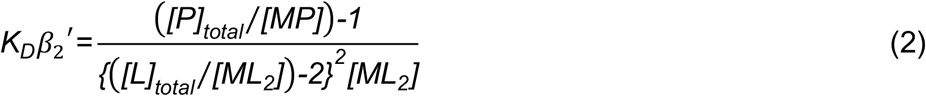

CopS_(34-151)_ - Cu^2+^ *K_D_* were determined using the indicator PAR as competitor ([Cu^II^(PAR)] conditional *K_A_’* formation constant for Cu^2+^ at pH 7.4 of 10^14.6^ M^−1^, isosbestic point *A*_445 nm_, *ε*_505 nm_ 41500 M^−1^cm^−1^(77)). 4 μM Cu^+^, 10 μM PAR in buffer 20 mM Hepes pH 7.4, 150 mM NaCl were titrated with 2 - 20 μM Strep-tagged CopS_(34-151)_, incubated at room temperature to equilibrate until no further spectral changes were observed (60 min), and the 300-800 nm absorption spectra recorded. *K_D_* value was obtained from a curve-fitting of a series of experimental data to equations (3) and (4). Reported errors are asymptotic standard errors provided by the fitting software (KaleidaGraph; Synergy).

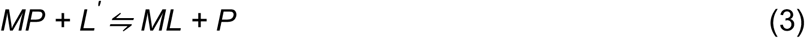

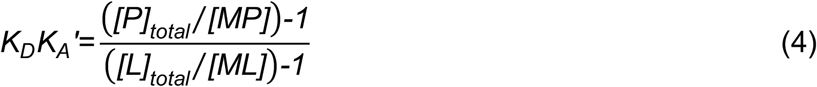

### Bioinformatic approaches

Protein sequences were retrieved from Uniprot (78), and aligned using Clustal Omega (79). To build the phylogenetic trees, the full-length protein sequences of *E. coli* CusS and *P. aeruginosa* CopS sequences were independently used as query to search for homologs in the UniProtKB database using the Uniprot/Blast tool. Sequences more than 45% identical over their entire lengths were retrieved and aligned. Phylogenetic trees were calculated with the Jalview software (80), using the distance matrix BLOSUM62 and the Average Distance (UPGMA) algorithm.

The structure of the soluble periplasmic copper binding loop of CopS_(34-151)_ was modeled using the online server SWISS-MODEL (81) and the structure of the *E. coli* CusS soluble periplasmic domain (PDB ID: 5KU5) (43) as template. Conserved metal binding residues of CopS were identified by superimposing its structure with 5KU5 using UCSF Chimera (82).

## Acknowledgments

This work was supported by grant R01GM114949 from the National Institutes of Health to JMA. FCS is a career investigator of CONICET and the Rosario National University Research Council.

## Footnotes

### Author contributions

LNA: performed research; LNA, FCS and JMA, designed research, analyzed data and wrote the paper.

The authors declare no competing interest.

## SUPPLEMENTAL INFORMATION

**Table S1.**
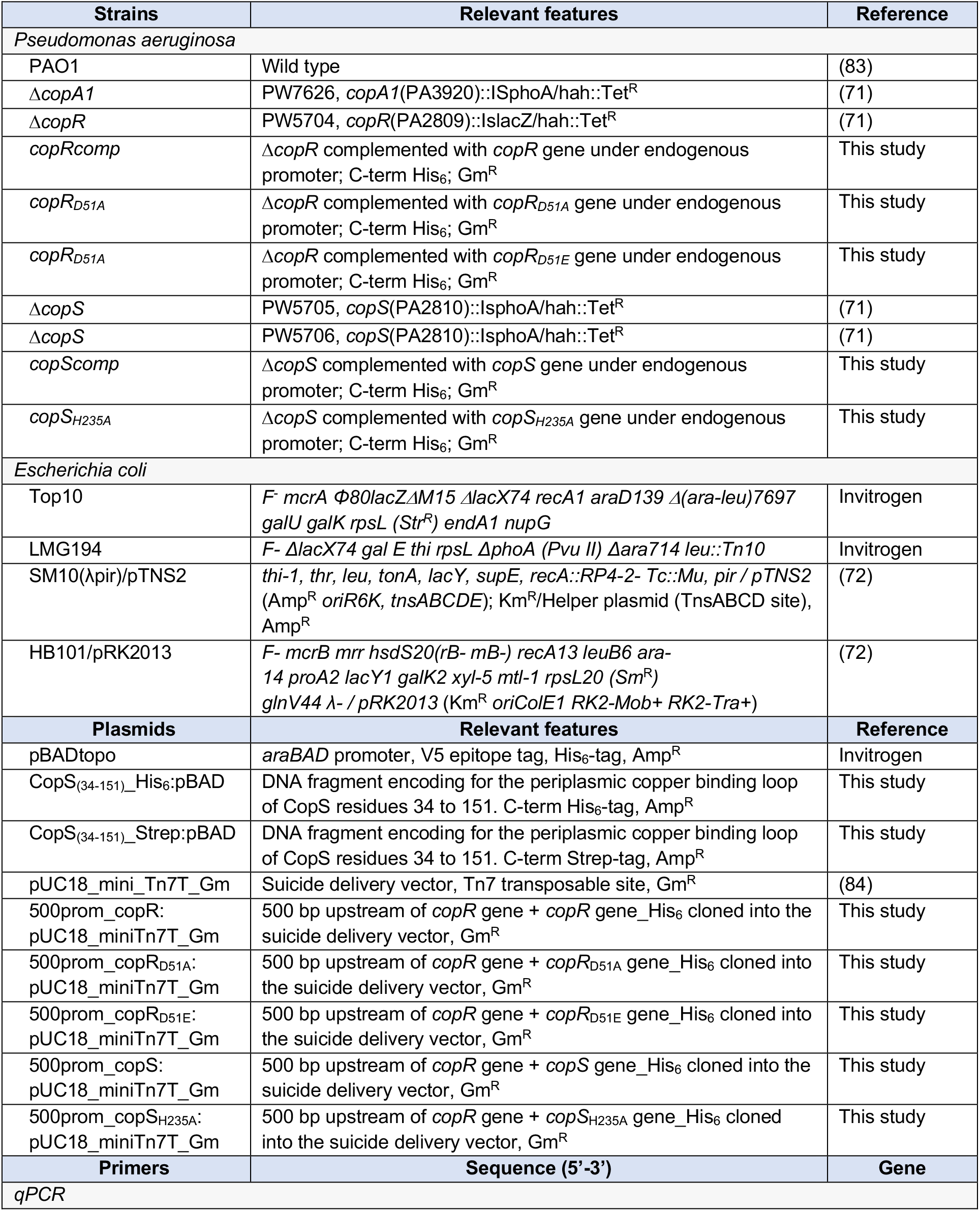
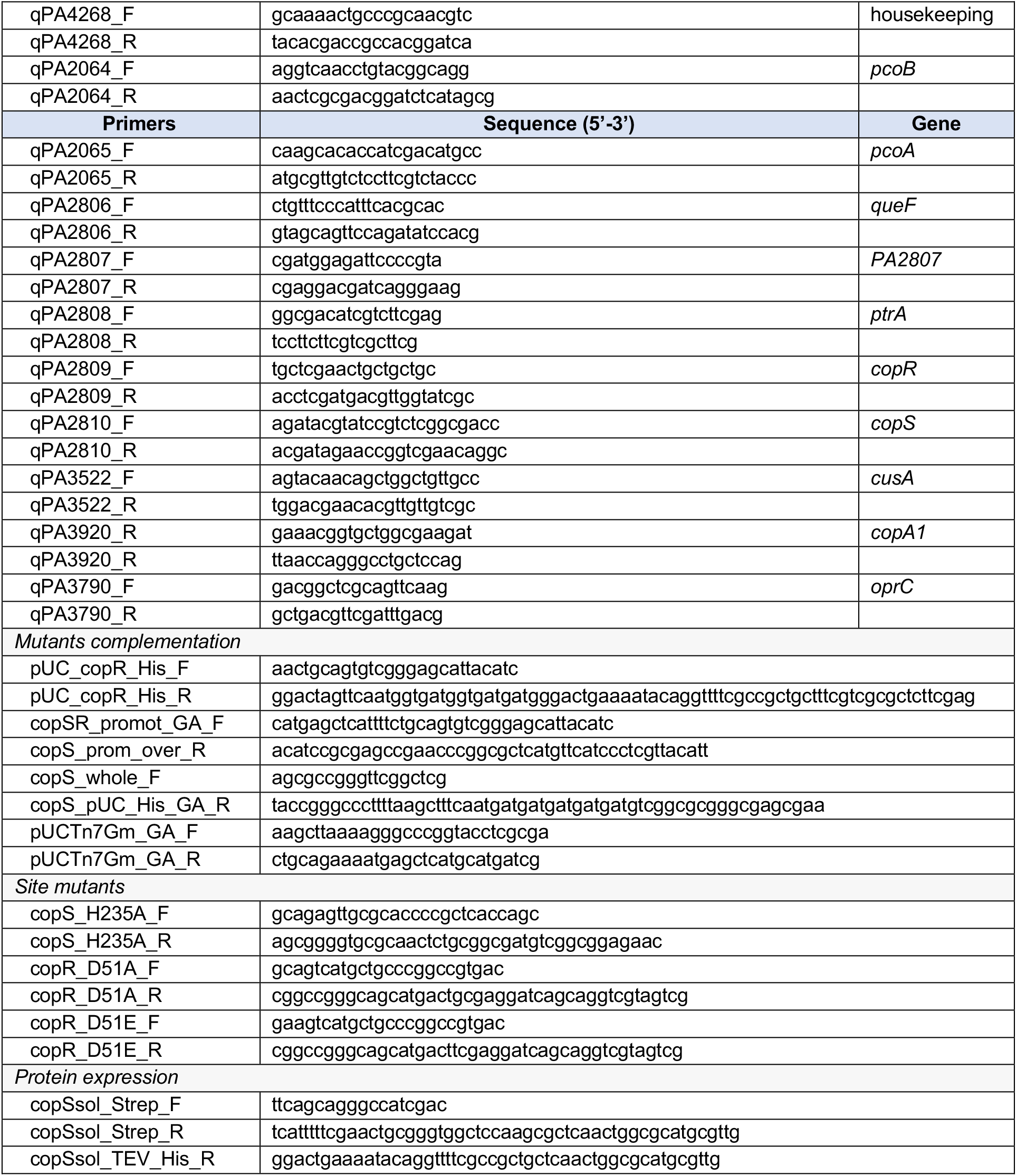
Bacterial strains, plasmids and primers used in this study.

---

**Fig. S1.**
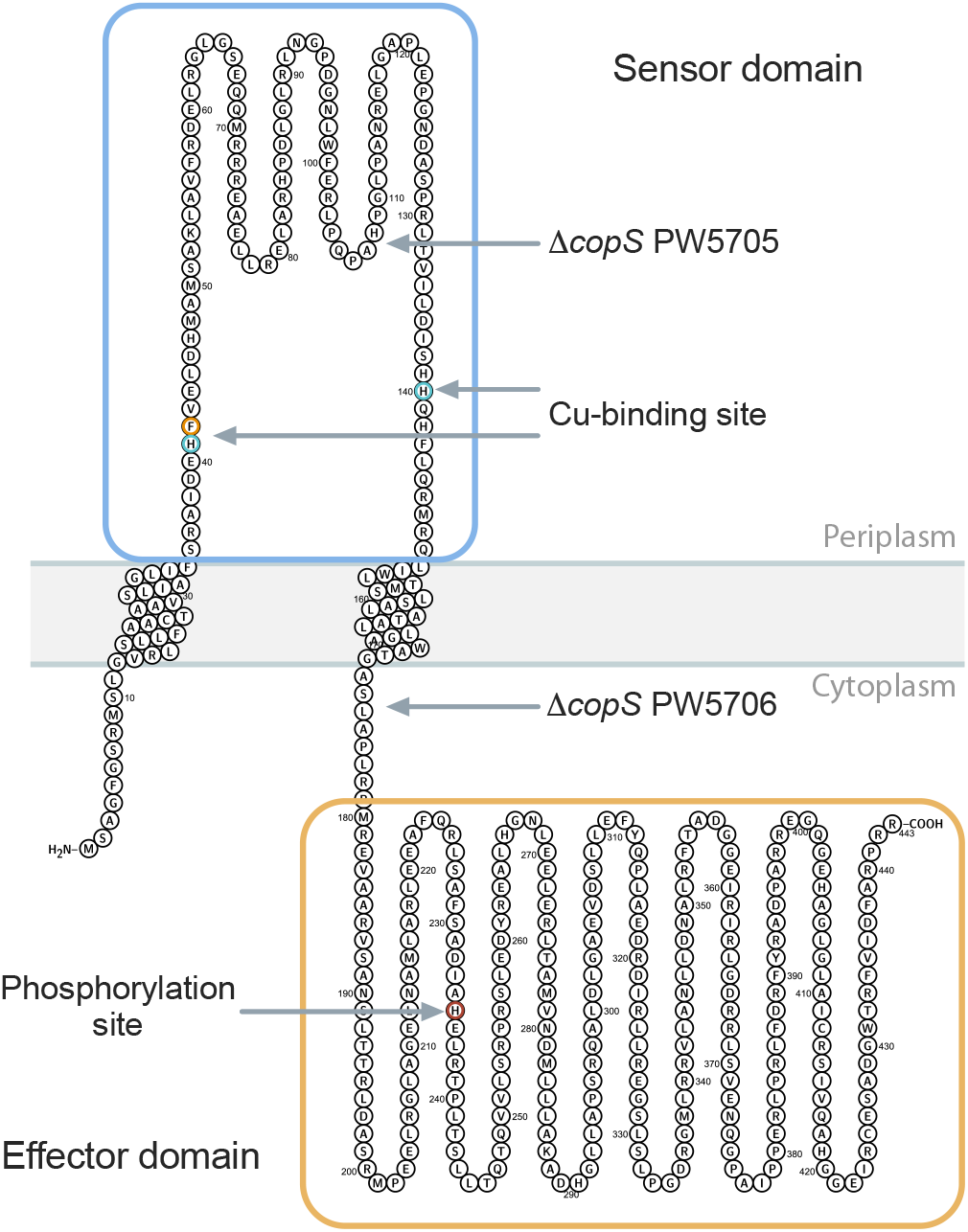
Topology, functional domains, and location of transposon insertions in CopS. The periplasmic Cu^+^ sensor domain of CopS is highlighted in blue. His_41_ and His_140_ in cyan, and Phe_42_, in orange are the residues forming the metal binding site. The C-terminal, cytoplasmic, effector domain (yellow), contains the phosphorylatable His_235_ (red). Both insertional mutants, PW5705 and PW5706, have an in-frame stop codons, producing shorter versions of CopS, lacking either part of the Cu-binding residues and the effector domain (PW5705), or just the effector domain (PW5706). CopS topology model was created using the Protter online tool version 1.0 (85).

**Fig. S2.**
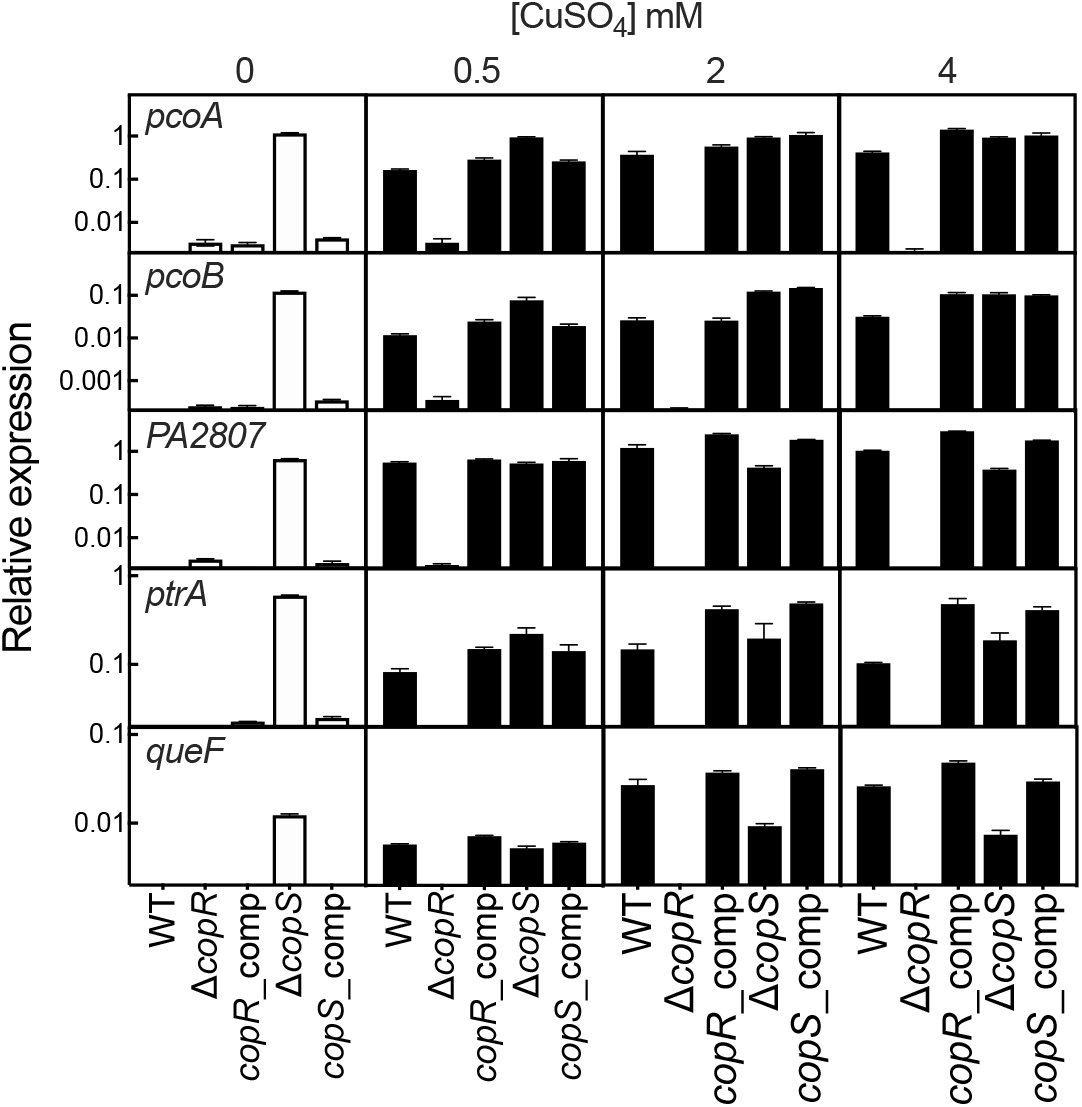
Expression of genes in the CopRS regulon in the Δ*copR* and Δ*copS* mutant strains quantified in the absence (white) and the presence (black) of 0.5, 2, and 4 mM CuSO_4_ (5 min treatment). Transcript levels of *pcoA*, *pcoB*, *PA2807*, *ptrA,* and *queF* genes are plotted relative to that of the housekeeping gene *PA4268*. Data are the mean ± SEM of three independent experiments.

**Fig. S3.**
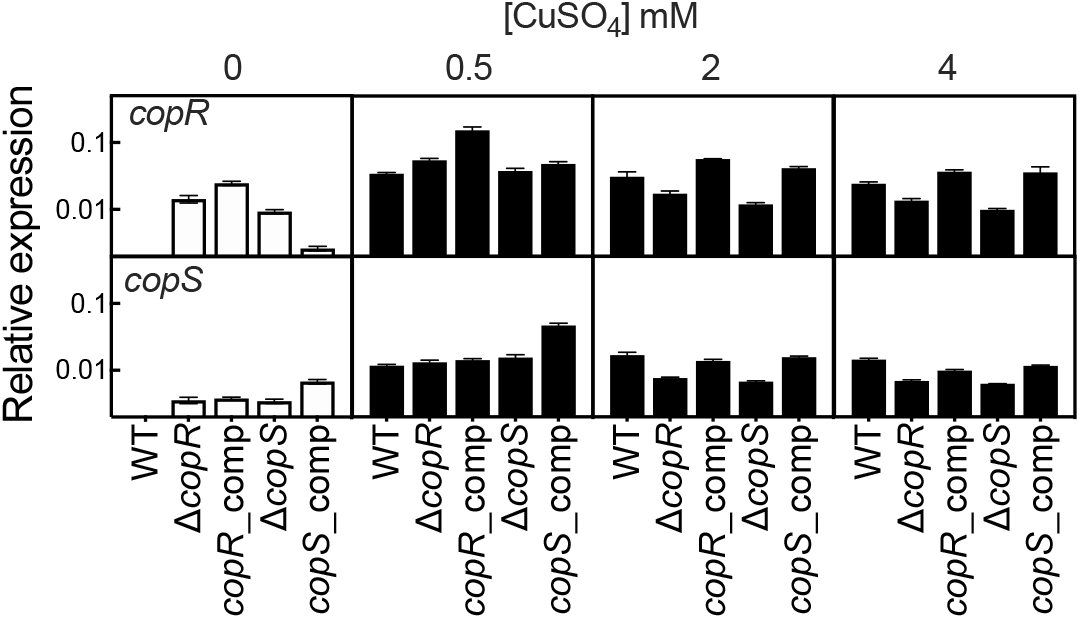
Expression of the *copRS operon* in the Δ*copR* and Δ*copS* mutant strains quantified in the absence (white) and the presence (black) of 0.5, 2, and 4 mM CuSO_4_ (5 min treatment). Transcript levels of *copR*, and *copS* genes are plotted relative to that of the housekeeping gene *PA4268*. Data are the mean ± SEM of three independent experiments.

**Fig. S4.**
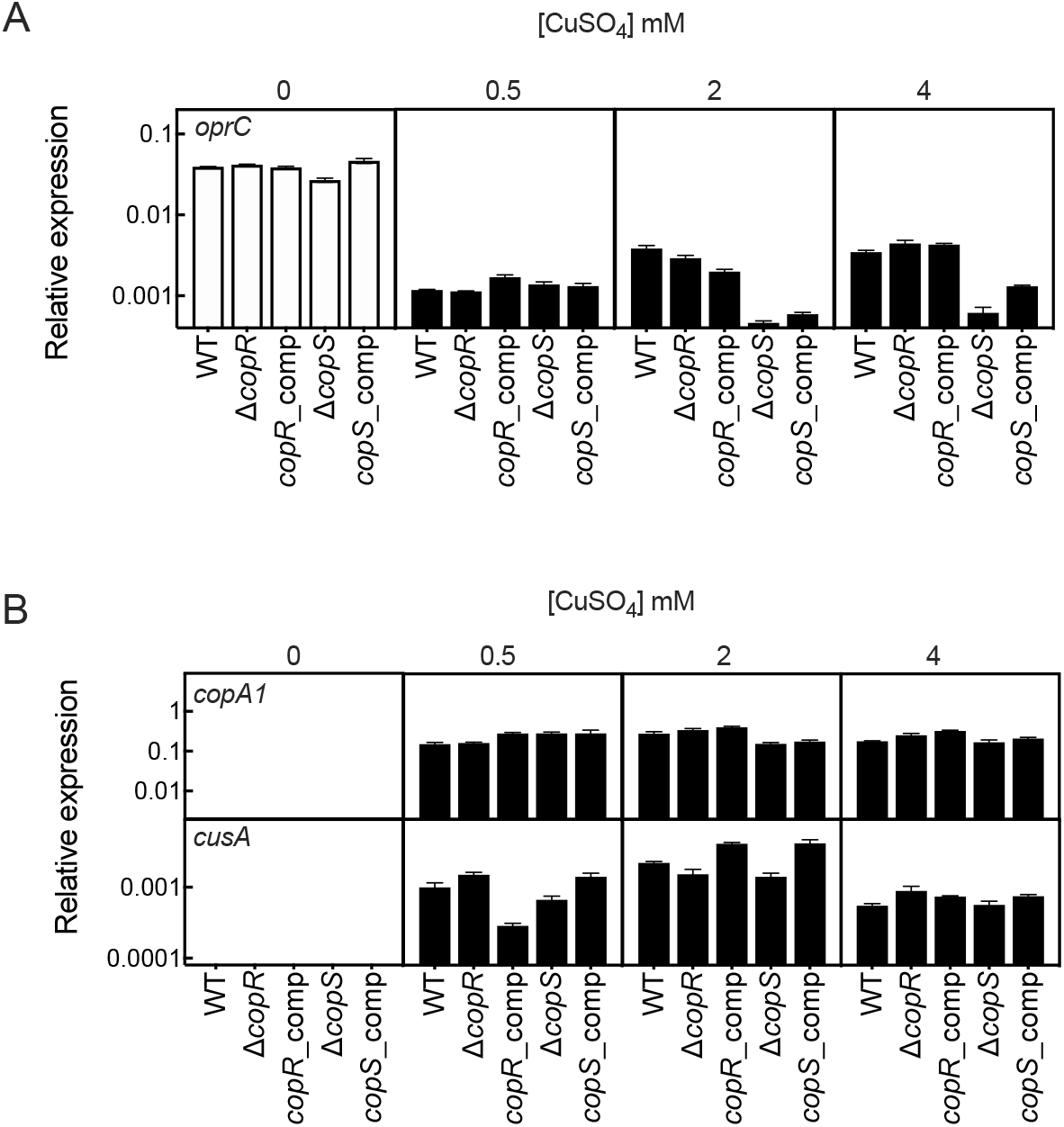
Expression of Cu transporter genes in the Δ*copR* and Δ*copS* mutant strains quantified in the absence (white) and the presence (black) of 0.5, 2 and 4 mM CuSO_4_ (5 min treatment). Transcript levels of *copA1,* coding for the Cu^+^ efflux P_1B_-type ATPase CopA1, *cusA,* a component of the RND CusABC system (A), and *oprC,* codding for Cu importer OprC (B), are plotted relative to that of the housekeeping gene *PA4268*. Data are the mean ± SEM of three independent experiments.

**Fig. S5.**
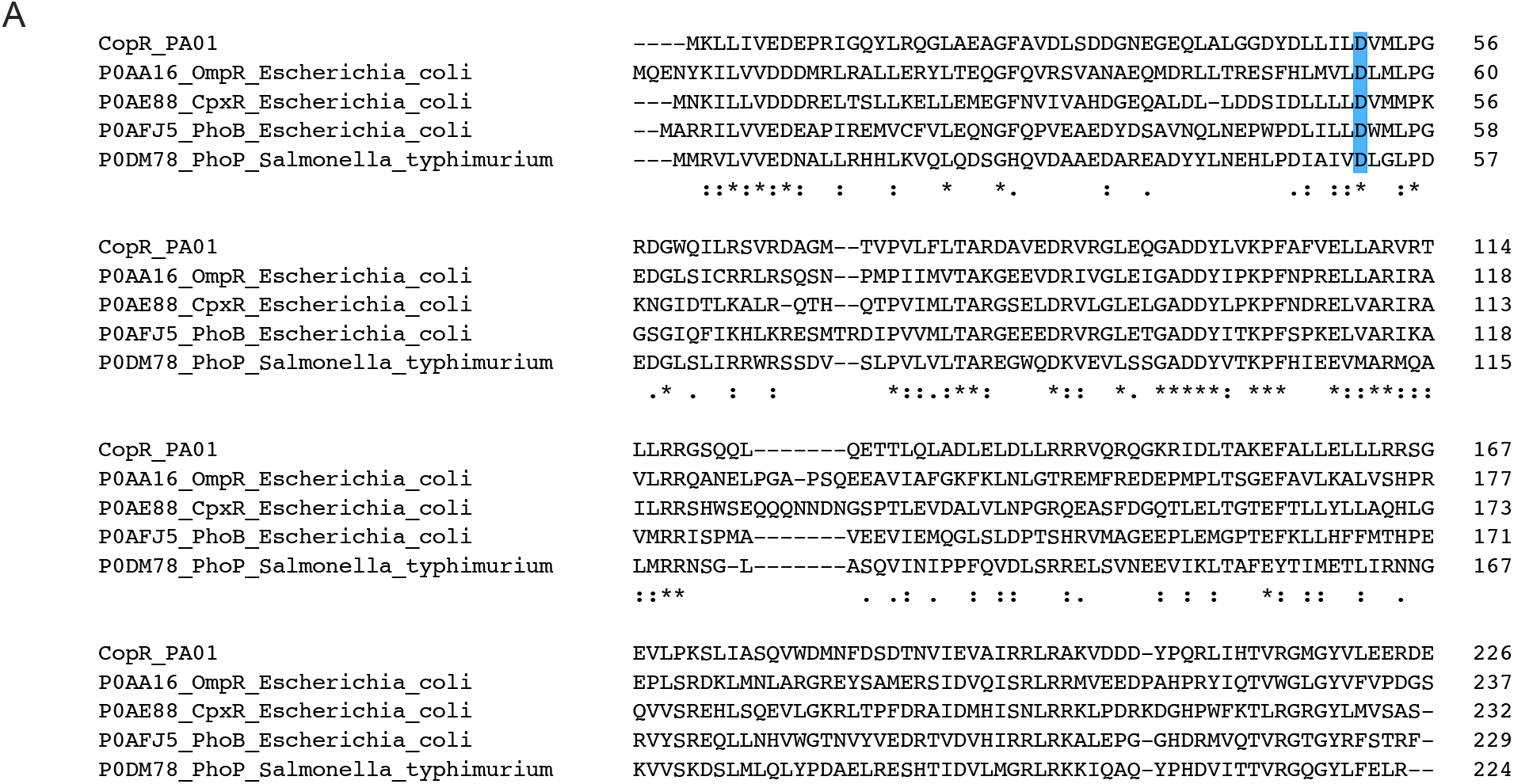
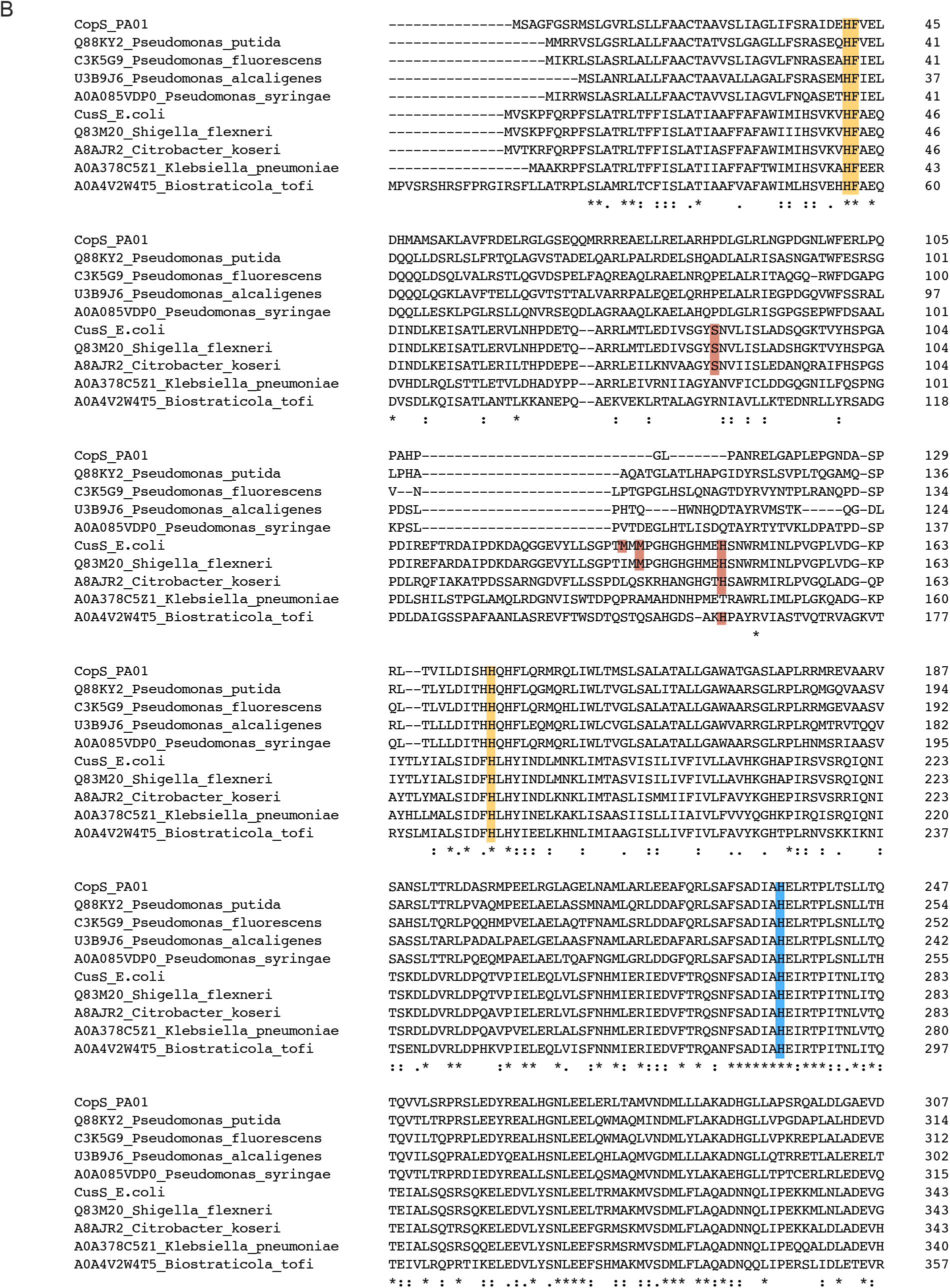
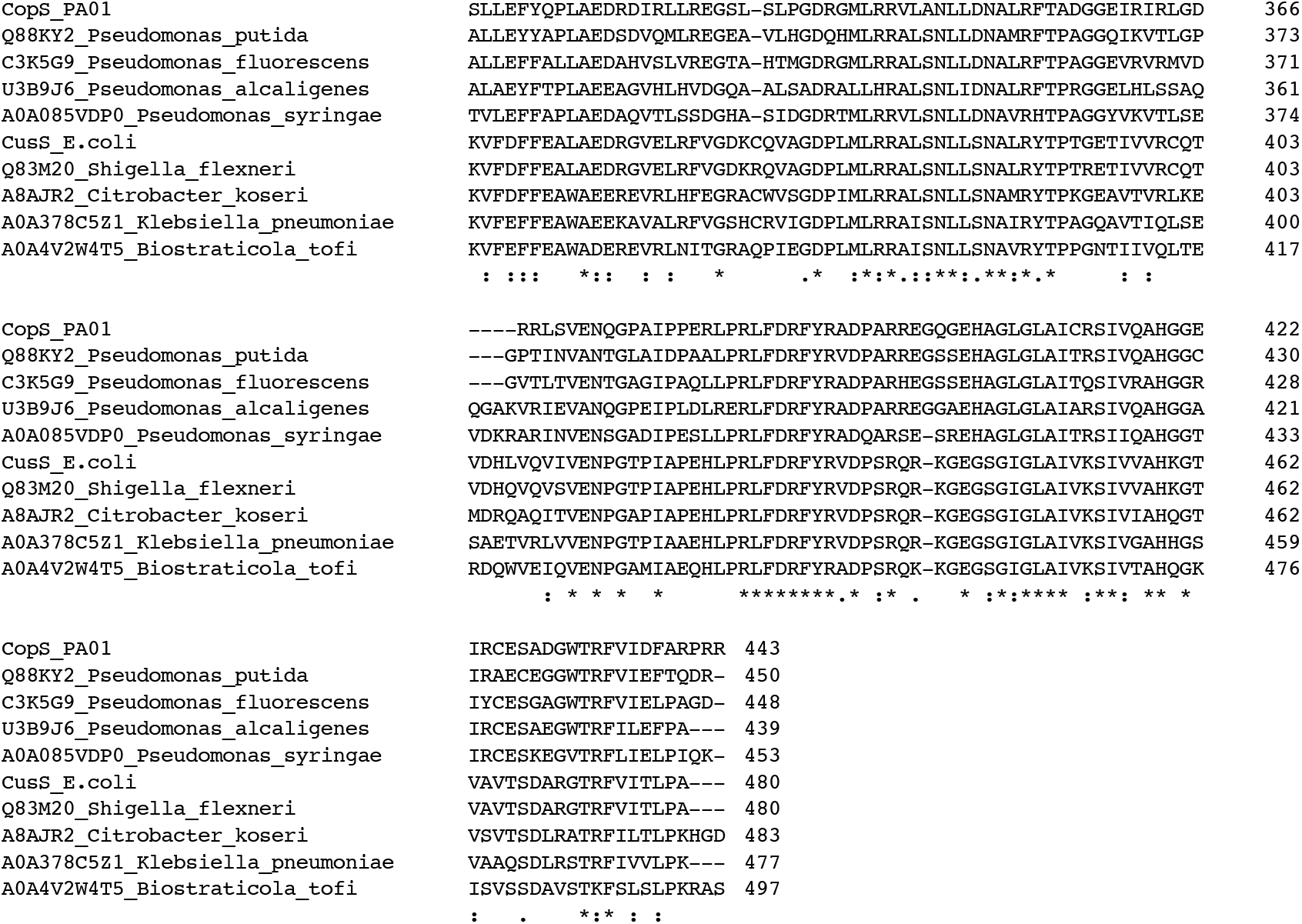
Multiple sequence alignment of the *P. aeruginosa* CopRS TCS proteins with bacterial homologs. (A) *P. aeruginosa* CopR protein sequence was aligned with characterized bacterial RR to identify the conserved phosphorylatable Asp residue (highlighted in blue). (B) *P. aeruginosa* CopS and *E. coli* CusS protein sequences were aligned with homologs of both of CopS-like and CusS-like proteins from different species. Conserved Cu binding sites at the dimeric interface are highlighted in yellow. *E. coli* CusS Cu binding sites, not conserved in CopS, are highlighted in orange. The conserved phosphorylatable His residue is highlighted in blue. Uniprot accession numbers precede each species name.

**Fig. S6.**
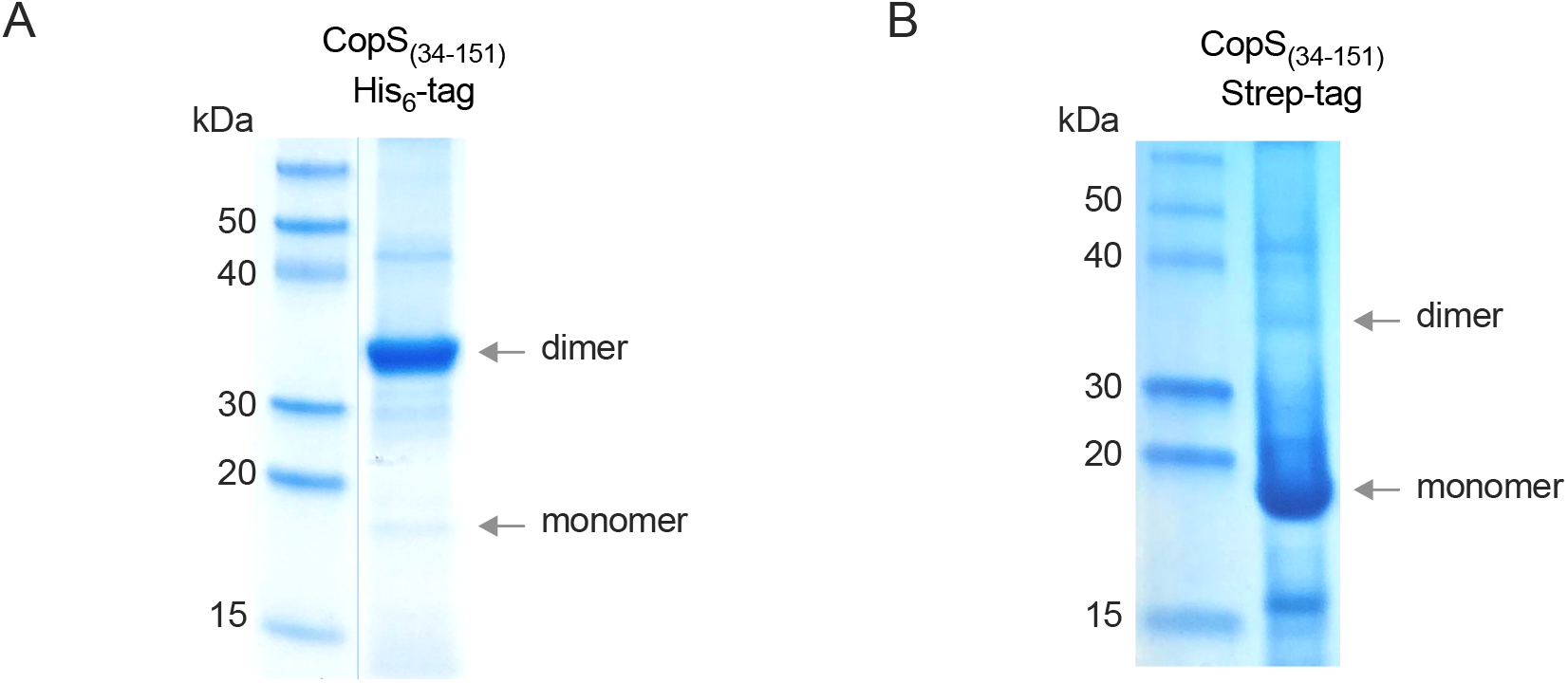
SDS-PAGE analysis of the periplasmic copper binding loop of CopS_(34-151)_. 10 μg of purified (A) His_6_-tagged or (B) strep-tagged protein were subjected to 8-to-16% gradient SDS-PAGE. Gels were stained with Coomassie Blue G250. Left lanes: molecular weight marker. Right lanes: purified proteins. Arrows indicate the protein monomers and dimers, with expected masses of 19 and 38 kDa, respectively. The presence of the C-terminal His_6_-tag in CopS_(34-151)_ stabilized the dimer form of the protein. The C-terminal Strep-tag did not. The gel shown in (A) was spliced for labeling purposes (blue vertical line).

**Fig. S7.**
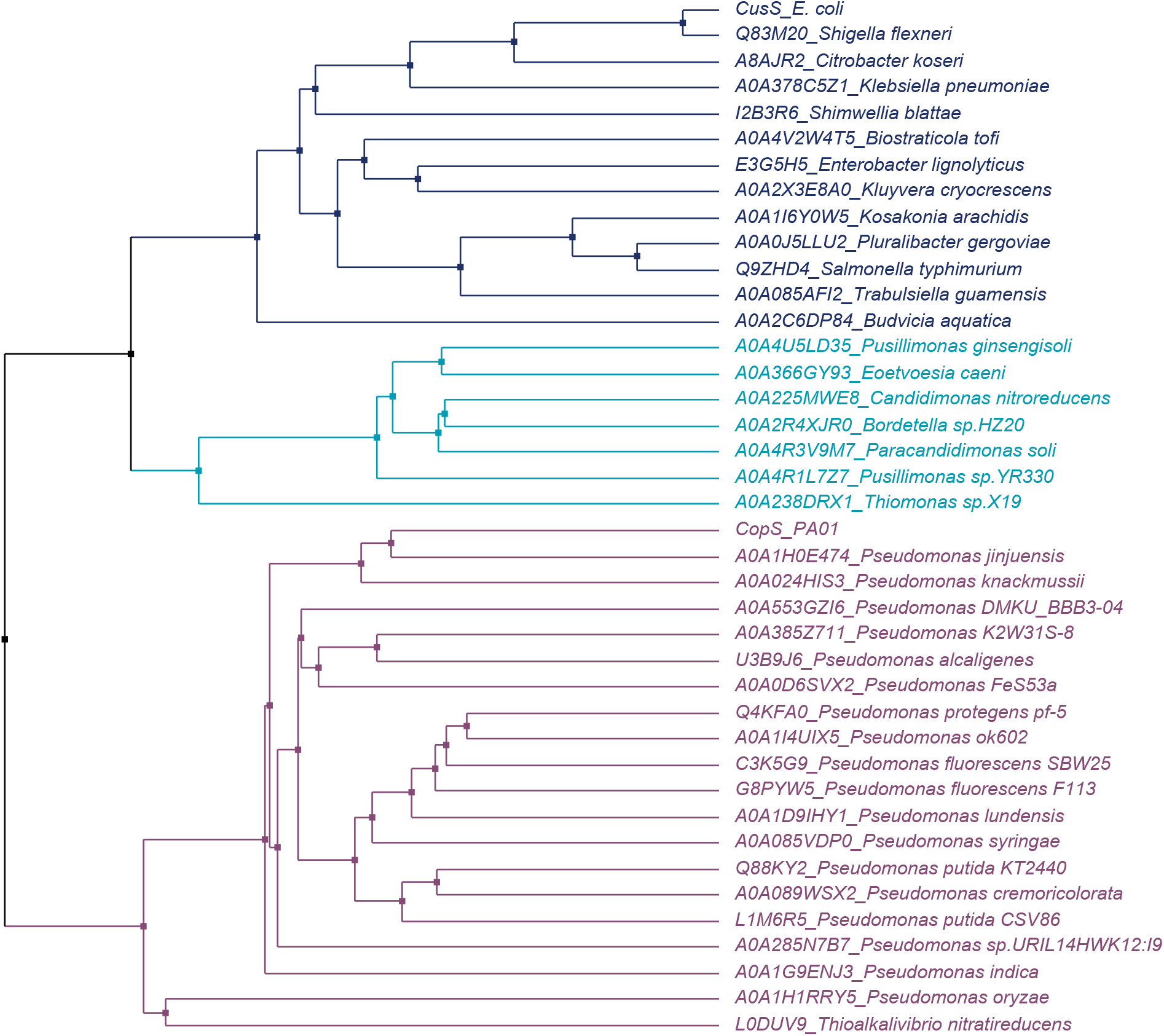
Phylogenetic tree of CopS-like and CusS-like proteins. Separately, *P. aeruginosa* CopS and *E. coli* CusS were used to find homologs in the UniProtKB database. The top-20 hits (>45% homology) from each Blast search were aligned with Clustal Omega, and the resulting alignment used to construct the displayed average distance tree. Different taxa were colored as follow: dark blue, Enterobacterales; cyan, Burkholderiales; and pale violet, Pseudomonadales. Uniprot accession numbers precede each species name.

